# Entropy of city street networks linked to future spatial navigation ability

**DOI:** 10.1101/2020.01.23.917211

**Authors:** A. Coutrot, E. Manley, S. Goodroe, C. Gahnstrom, G. Filomena, D. Yesiltepe, R.C. Dalton, J. M. Wiener, C. Hölscher, M. Hornberger, H. J. Spiers

## Abstract

Cultural and geographical properties of the environment have been shown to deeply influence cognition and mental health[1–6]. While living near green spaces has been found to be strongly beneficial [7–11], urban residence has been associated with a higher risk of some psychiatric disorders [12–14] (although see [15]). However, how the environment one grew up in impacts later cognitive abilities remains poorly understood. Here, we used a cognitive task embedded in a video game[16] to measure non-verbal spatial navigation ability in 397,162 people from 38 countries across the world. Overall, we found that people who grew up outside cities are better at navigation. More specifically, people were better at navigating in environments topologically similar to where they grew up. Growing up in cities with low Street Network Entropy (e.g. Chicago) led to better results at video game levels with a regular layout, while growing up outside cities or in cities with higher Street Network Entropy (e.g. Prague) led to better results at more entropic video game levels. This evidences the impact of the environment on human cognition on a global scale, and highlights the importance of urban design on human cognition and brain function.

## Introduction

Cognitive abilities, including spatial navigation, have been shown to correlate with specific genotypes [17]. However, research on brain plasticity supports the notion that experience shapes brain structure as well as function [2, 3]. In particular, cultural and geographical properties of the environment have been shown to deeply influence cognition and mental health [4, 6]. In rodents, exploring complex environments has a positive impact on hippocampal neurogenesis and cognition [1, 5]. In humans, spatial navigation activates the hippocampus [18], and continuous navigation of a large complex city environment increases posterior hippocampal volume [19]. However, how the environment one grew up in impacts later cognitive abilities remains poorly understood for two reasons. First, human environments are manifold and much harder to characterize than a rodent’s cage. Second, collecting cognitive data of large samples from populations living in different environments is very costly [20]. To overcome these limitations, we measured non-verbal spatial navigation ability in 3.9 million people across all countries and examined a subset of this data (397,162 people, in 38 countries). We used a cognitive task embedded in a video game, that is predictive of real-world navigation skill [16, 21, 22]. While the task has been described previously [16], we now report new data not previously published. We focused on spatial navigation due to its universal requirement across cultures, and parallels to rodent studies [23, 24]. We quantified the complexity of participants’ environments with OSMnx, a tool giving access to the street network topology of cities anywhere in the world [25]. We found that, on average, people who reported having grown up in cities have worse navigation skills than those who reported growing up outside cities, even when controlling for age, gender, and level of education. This difference between city and non-city people varied across countries, being – for instance – more than 6 times larger in the USA than in Romania. To investigate these variations we computed the average Street Network Entropy (SNE) of the biggest cities of 38 countries; grid-like cities (e.g. Chicago) have a small SNE, while more organic cities (e.g. Prague) have a higher one. We found that growing up in cities with low SNE led to better performance at video game levels with a regular layout, while growing up outside cities or in cities with higher SNE led to better performance at more entropic video game levels. This confirms the impact of the environment on human cognition on a global scale, and highlights the importance of urban design on human cognition and brain function.

## Results and discussion

We used the Sea Hero Quest database, which contains the spatial navigation behaviour of 3.9 million participants measured with a mobile video game, ‘Sea Hero Quest’ (SHQ) [16, 22]. SHQ involves navigating a boat in search of sea creatures (Figure 1). Performance of SHQ has been shown to be predictive of real-world navigation ability [21]. It has also allowed to differentiate high-risk preclinical Alzheimer’s disease cases from low-risk participants [26]. Here, we focused on the wayfinding task [16], where players are initially presented with a map indicating the start location and the location of several checkpoints to find in a set order. To provide a reliable estimate of spatial navigation ability, we examined the data only from participants who had completed a minimum of eleven levels of the game, and who entered all their demographics (see Methods). This resulted in 397,162 participants from 38 countries included in our analysis, (see Table S1). Among them, 212,143 males (mean age: 37.81 ± 13.59 years old) and 185,173 females (mean age: 38.67 ± 14.92 years old).

**Figure 1.**
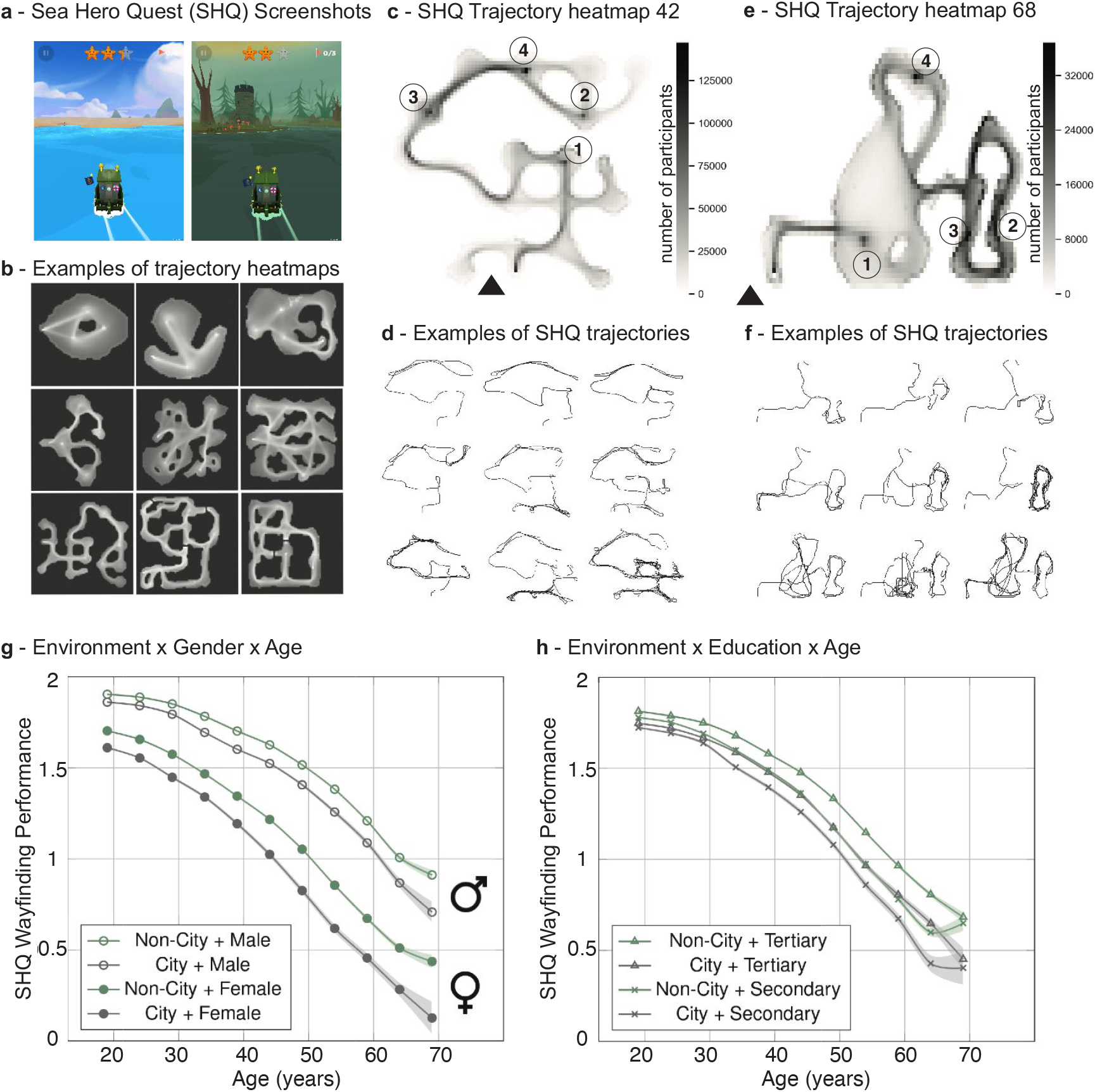
Wayfinding task. **a** Screenshots from the game Sea Hero Quest (SHQ). See also Supplementary Video 1. **b** Nine examples of trajectory heatmaps out of the 75 SHQ levels. **c - e** Heatmaps of the trajectories of all participants in level 42 (N=171,887) and level 68 (N=40,251) of SHQ.The black triangle represents the starting position, and the circled numbers represent the ordered checkpoints the participants must reach. **d - f** Examples of trajectories in level 42 and 68 of SHQ. **g - h** - Association between Environment and SHQ Wayfinding Performance stratified by age, gender, and education. The SHQ Wayfinding Performance is computed from the trajectory length and has been averaged within 5-years windows. See Extended Data Fig. 2 for a breakdown of the environment classes. The error bars represent the standard errors and the center values correspond to the means.

### The association between environment and spatial ability stratified by age, gender and education

To quantify spatial ability, we defined the “wayfinding performance” metric (*WF*), which captures how efficient participants were in completing the wayfinding levels, while correcting for video-gaming skills (see Methods). We performed the same analysis for path integration levels, see Supplementary Methods and Extended Data Fig. 6. It yielded similar results. A multivariate linear regression was calculated to predict wayfinding performance based on age, gender, education and environment. Age has the strongest effect (*F*_1,397157_ = 61389, *p* < 0.001, *η*^2^ = 0.127, Hedge’s *g* = 0.98, 95% CI = [0.97, 0.99]), followed by gender (*F*_1,397157_ = 20665, *p* < 0.001, *η*^2^ = 0.043, Hedge’s *g* = 0.44, 95% CI = [0.43, 0.45]), education (*F*_1,397157_ = 1476.9, *p* < 0.001, *η*^2^ = 0.003, Hedge’s *g* = 0.13, 95% CI = [0.13, 0.14]), and environment (*F*_1,397157_ = 1628.8, *p* < 0.001, *η*^2^ = 0.003, Hedge’s *g* = 0.09, 95% CI = [0.09, 0.10]). The Hedge’s g of age is computed between participants under 25 years old (N=88,101) and above 55 years old (N=59,982). Figures 1g-h represent the effect of the environment on *W F* stratified by age, gender, and education. We replicate previous studies showing that wayfinding performance decreases with age [27, 28], males perform better than females [29], and performance increases with the level of education [30, 31]. Here we now report that participants raised outside cities are more accurate navigators than city-dwellers. Having a tertiary level of education while having grown up in a city is roughly equivalent to having a secondary level of education while having grown up outside cities in terms of wayfinding performance. The sample sizes for each demographic and country are available Table S1. Given the magnitude of the dataset, most effects are likely to always be ‘significant below the 0.001 threshold’. In the following, we will focus on effect sizes as they are independent of sample size. We computed Hedge’s g between the city and not-city groups. To marginalize the effect of age, we computed Hedge’s g within 5-years windows, see Extended Data Fig. 1. Averaged over all age groups, g = 0.13, 95%CI=[0.12, 0.14], positive values corresponding to an advantage for participants who grew up outside cities. As shown in Figure S1, Hedge’s g remained stable across age. This stability is interesting as one could have hypothesized that the influence of the environment one grew up in could fade with age. This stability is consistent with the literature on the timing of enriched environment exposure in mice, showing that an early enriched environment exposure could provide a “reserve”-like advantage which supports an enduring preservation of spatial capabilities in older age [32].

### The association between environment and spatial ability across countries

To quantify how spatial ability and environment are associated across countries, we fit a Linear Mixed Model (LMM) for wayfinding performance, with fixed effects for age, gender and education, and a random effect for country, with random slopes for environment clustered by country: *WF*_*perf*_ ∼ *age* + *gender* + *education* + (1 + *environment/country*). Figure 2a represents the environment slopes for each country, positive values indicate an advantage for participants raised outside cities. In terms of Hedge’s g, this spectrum ranges from Romania (g = -0.03, 95%CI=[-0.10, 0.04])) to the United States (g = 0.19, 95%CI=[0.17, 0.21]). Figure S2 illustrates this difference in effect size across age in different countries. For instance, while the effect size is close to being null in Germany, growing up in cities in the USA cost the equivalent of aging four years in terms of spatial ability, see Extended Data Fig. 3.

**Figure 2.**
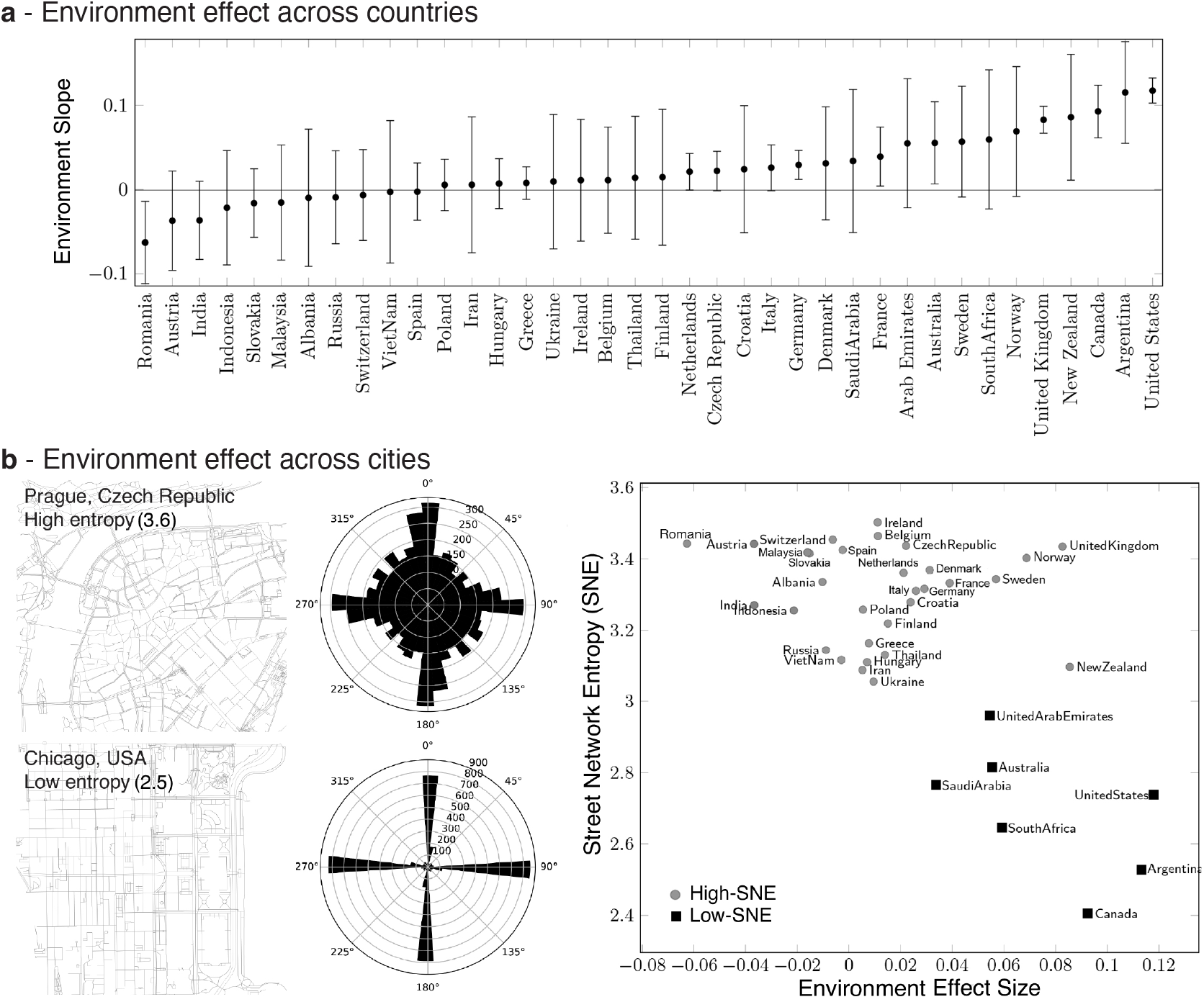
Street Network Entropy (SNE) and environment effect in 38 countries. **a** - Differences across countries. We fit a linear mixed model for wayfinding performance, with fixed effects for age, gender and education, and random environment slopes clustered by country (n=397,162 participants). We plot the environment effect sizes (country slopes) for each country, with positive values indicating an advantage for participants raised outside cities. Error bars correspond to standard errors. **b** - Left: Two examples cities with low (Chicago, USA) and high (Prague, Czech Republic) SNE. See also Extended Data Fig. 10. Distribution of the street bearings across 36 bins of 10 degrees. Right: Average SNE as a function of the environment effect size (random environment slope) in each country. Positive values indicate an advantage of participants raised outside cities compared to their urban compatriots. Average SNE is the weighted average over the 10 most populated cities of the country, weighted by their population. Squares and circles correspond to the low-SNE and high SNE country groups, determined with k-means.

### Associations between the topology of cities and spatial ability

To explain the variations in the association between environment and spatial ability across countries, we hypothesized that countries with lower effect sizes contain cities with more complex layouts, which places greater demands on navigation, honing the skill of those growing up in them. The impact of city topology on human spatial ability has previously been theorized in the urban design literature [33, 34], but the empirical studies on street networks suffer from limitations, mostly due to data availability, gathering, and processing constraints [35, 36], although see [37]. To overcome these limitations, we coupled our global dataset with the OSMnx toolbox, which provides the street network layout for anywhere in the world via OpenStreetMap [25, 38]. Street network complexity is a manifold concept, and many metrics have been proposed to quantify it. Shannon’s information entropy [39] is arguably the simplest and the most general measure of network complexity, from neural to spatial networks [40–43]. The entropy of a variable can be interpreted as the average level of uncertainty inherent in its possible outcomes. Here, we computed the Shannon entropy of the city’s street orientations’ distribution. The smaller the entropy, the less complex - i.e. the more ordered - the city street network, see examples in Figure 2b and in Extended Data Fig. 10. Since SHQ participants only reported their home country, and not finer-grained regional information, we computed the Street Network Entropy (SNE), defined as the average of the entropy of the ten biggest cities of each country in terms of population, weighted by the city population (Table S2). Thus, we had one SNE value per country. Figure 2b represents the SNE of countries as a function of their environment slope from the above mixed model. The majority of the countries have a similar SNE, corresponding to the typical organic street pattern of old city cores (e.g. France, Romania, Spain, but also Thailand or India). However some other countries have distinctly smaller SNE, corresponding to the orthogonal grid, a very common planned street pattern (e.g. the United States, Argentina). The bivariate correlation between country SNE and their environment slope is significant (Pearson’s *r*(36) = −0.60, *p* < 0.001, 95%CI =[-0.78 -0.30]). This validates our hypothesis: the lower the SNE (i.e. the simpler the street network), the worse the spatial ability of the people who grew up in cities compared to their compatriots raised outside cities. This effect remained when we control for Gross Domestic Product (GDP) per capita (linear regression predicting environment slopes, GDP per capita t(35)=4.02, *p* < 0.001, SNE t(35)=-5.86, *p* < 0.001), see Table 2 in Methods. We did not find a correlation between GDP per capita and SNE (r(36)=0.14, p=0.40), see Extended Data Fig. 7a.

While SNE captures the spatial organisation of a city, metrics based on the graph theoretic network measure topological properties of the cities, which could also play a role in shaping navigation skill. Across our 380 cities we measured the betweenness, closeness and degree centrality, average circuity, average neighborhood degree, clustering coefficient and average street length (see Methods) [43, 44]. After applying Bonferroni correction for multiple comparisons, we found that only circuity (network distance/Euclidean distance) was significantly correlated with SNE (r(36)=0.47, *p* = 0.02), which is coherent with [43]. This indicates that less regular city layouts are associated with paths across them that require more deviation around obstacles, see Figure 3.

**Figure 3.**
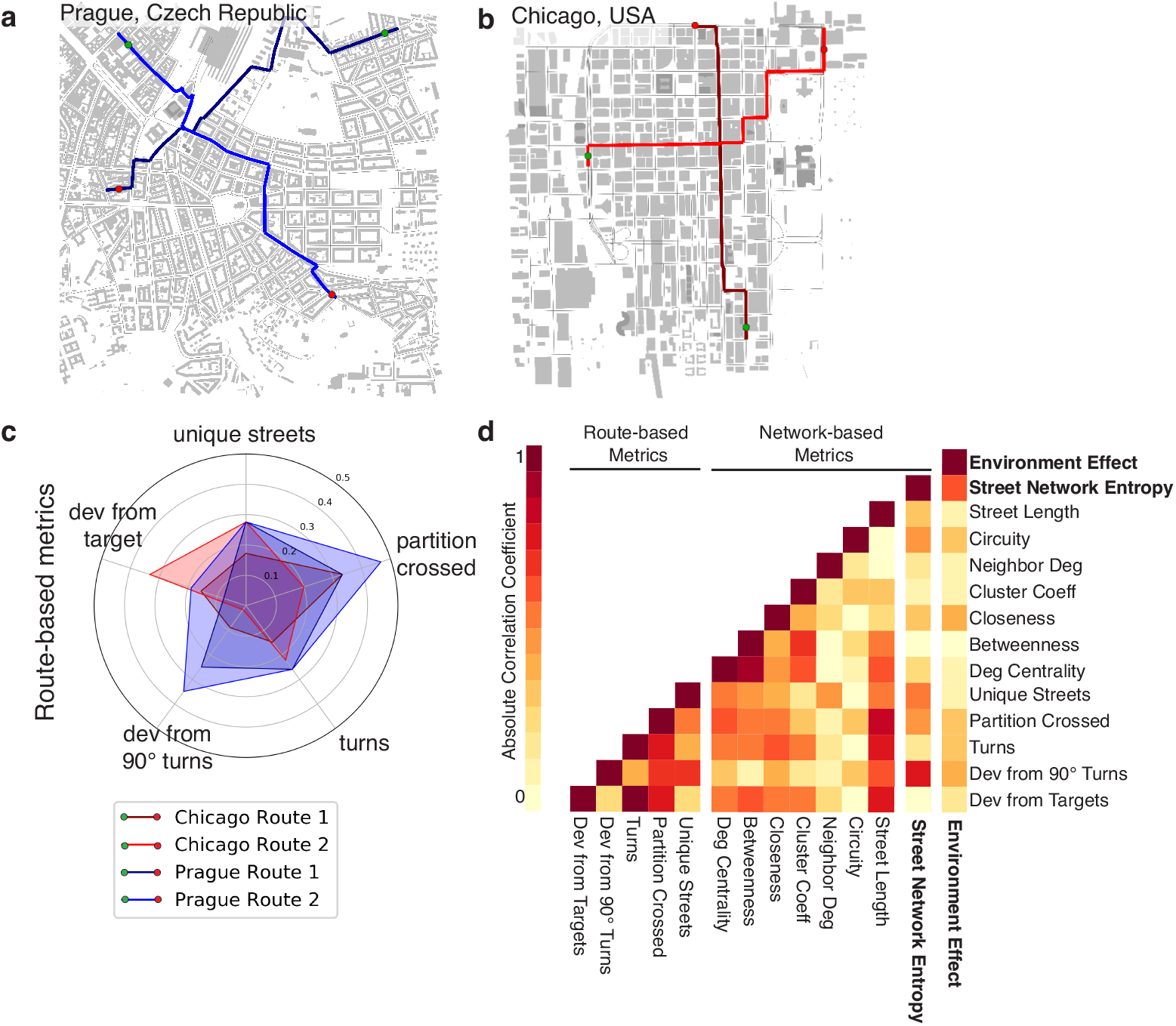
Comparison of Street Network Entropy (SNE) to other measures of city complexity measures. **a - b** We simulated 1000 routes in each of the 380 included cities. Four example representative routes in two contrasting cites high/low SNE are displayed. **c** We derived five key variables from each route: Number of above 50 degree turns, Number of unique streets, Deviation from regular 90 degree turns at each turn, Overall deviation from the target and Number of transitions in the partitions in street network structure. The spider plot shows the route variables for these 4 routes, for full visualisation of the average of the 1000 routes in the 380 included cities, see Extended Data Fig. 9. We also explored a range of other graph-theoretic measures commonly considered for spatial analysis of cities. **d** Absolute correlation coefficient between all the metrics and the environment effect size. SNE is by far the metric most correlated to the environment effect size.

What are the mechanisms by which exposure to high SNE would lead to better navigation ability? We surmised that navigating cities with irregular street layouts would require increased demands on: 1) Keeping track of the goal direction due to greater varying street angles; 2) Spatial/prospective memory for street names and upcoming turns due the human tendency to minimise the streets/turns in an irregular laid out city [45, 46]; 3) More hierarchical planning across neighbourhood, due to larger number of neighborhoods that might occur in irregular cities. Such demands would likely enhance the capacity of neural systems underlying orientation, prospective memory and planning abilities [19, 47]. To empirically determine whether the specific variables we propose are linked to SNE, we employed agent-based modeling to simulate 1000 routes through each of the 380 cities to quantify: number of turns, number of streets, deviation from a 90 degree turns, overall deviation from the target and number of crossed partitions in street network (see Methods and Figure 3). After applying Bonferroni correction for multiple comparisons, we found that turns and the deviation from the goal were not significantly higher in high-SNE cities, indicating that these may not be key factors in enhancing navigation skill. Rather we found that SNE was significantly correlated with deviation from 90 degree turns (r(36)=0.77, *p* < 0.001), number of streets (r(36)=0.57, *p* < 0.001), and number of partitions crossed (r(36)=0.48, *p* = 0.007). Thus, it appears that having to accommodate turns that deviate from 90 degrees and to navigate more streets and neighborhoods are key to enhancing navigation skill. We incorporated all network and route measures into a linear model to predict the environment effect size, and failed to find a significant equation (F(10,25)=1.44, *p* = 0.21). On the other hand, when using SNE only as a predictor, a significant equation was found (F(1,36)=20.1, *p* < 0.001), see Methods. This implies that it is the combination of navigational challenges that high SNE cities provide that is important for enhancing the inhabitants navigational skill.

### The interaction between virtual and real-world topologies and their influence on spatial abilities

We tested the symmetrical effect: do different SHQ level topologies interact with the effect of participant’s home environment? Here our hypothesis was that people growing up in environments with more complex topologies might perform better at more elaborate, entropic SHQ levels. Conversely, people growing up in regular cities might perform better at regular SHQ levels. We used k-means to split the countries into two SNE groups, revealing a low-SNE group and high-SNE group, see Figure 2b. We defined the SHQ level entropy as we did for the cities, with the orientations’ distribution computed from the level’s simplified Voronoi map (see Figure 4a and Methods for details). In order to include in our analysis as many level topologies as possible, we ran the following analysis on a subset of included participants who completed all SHQ levels (75 levels, 9,439 participants). We fit two LMMs for wayfinding performance: one with the participants from low-SNE countries (N=2021), the other with the participants from high-SNE countries (N=7418). Both models had fixed effects for age, gender and education, and a random effect for level, with random slopes for environment clustered by level: *WF*_*perf*_ ∼ *age* + *gender* + *education* + (1 + *environment/level*). We included all the wayfinding levels (N=42). Figure 4b represents for each level its entropy as a function of the environment slope *s*_*low*_ computed from participants in the low-SNE countries, and *s*_*high*_ computed from the high-SNE countries. Positive values correspond to an advantage of participants growing up outside cities. We observed that the only negative environment slopes correspond to low-SNE participants in less entropic levels, suggesting that people used to less entropic environments perform better in less entropic SHQ levels. The correlation coefficient was higher between the entropy of the levels and the low-SNE environment slopes (Pearson’s correlation *r*_*low*_ = 0.57, *p* < 0.001) than between the entropy of the levels and the high-SNE environment slopes (Pearson’s correlation *r*_*high*_ = 0.44, *p* = 0.003). This correlation slope difference did not reach statistical significance (Fisher’s z = 0.77, *p* = 0.22, 95% CI for *r*_*high*_ − *r*_*low*_ = [−0.46, 0.20]).

**Figure 4.**
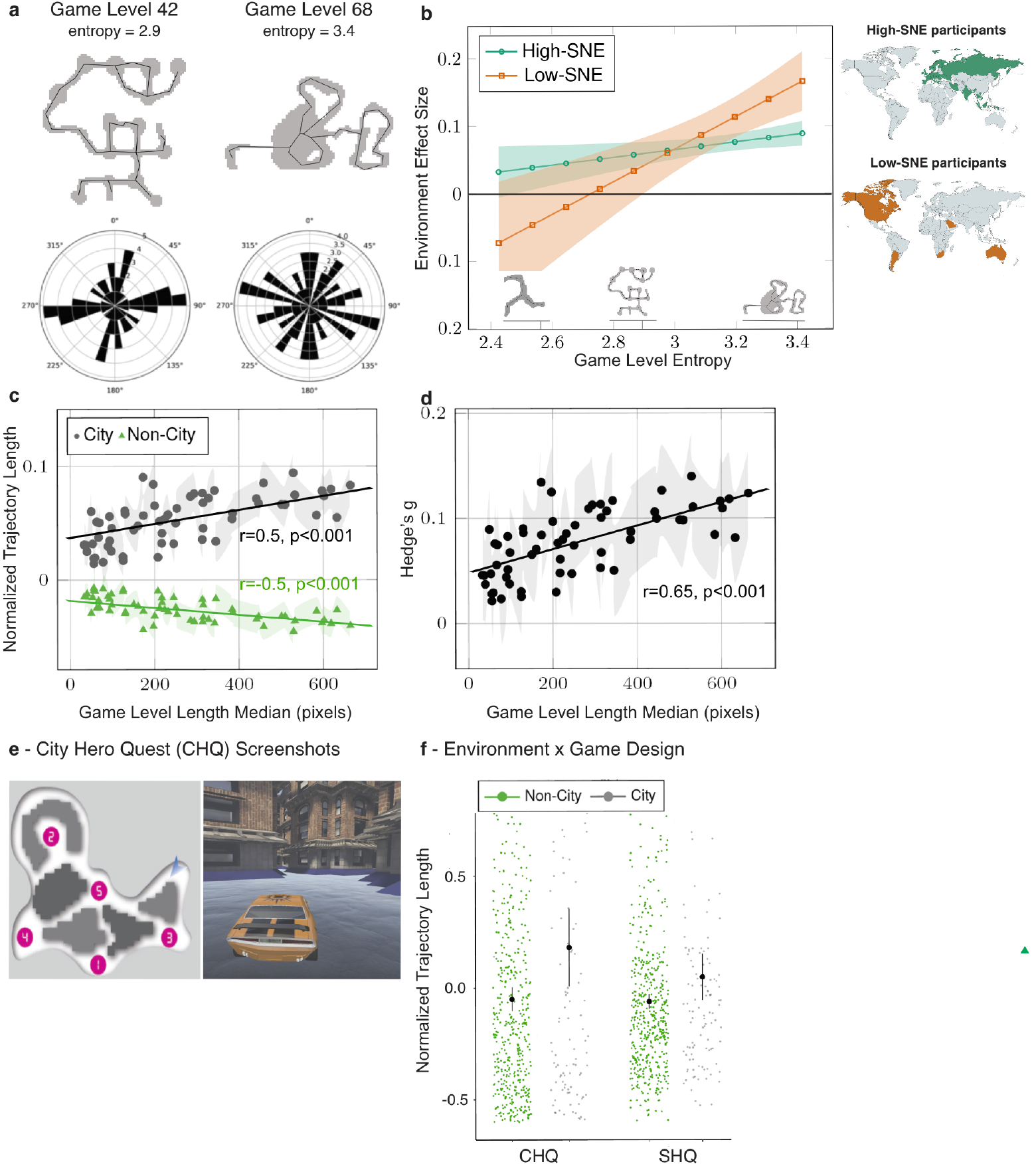
Participants are accurate at navigating more entropic game levels when they grew up in more entropic environments. **a** The entropy of the Sea Hero Quest (SHQ) levels is computed from the bearing distribution (rose plot), shown for 2 example levels. **b** Least square regression lines of the environment effect size on game level entropy for the High-SNE and low-SNE country groups (see mini-maps and Figure 2b). We included the players that played all the SHQ wayfinding levels (N = 10,626). Positive values indicate an advantage of participants raised outside cities. Low-SNE environment slopes are negative for low entropy SHQ levels, suggesting that in less entropic SHQ levels, people who grew up in less entropic cities perform better than their compatriots who grew up outside cities. **c** Normalized trajectory length as a function of the level length. Trajectory lengths have been z-scored for each level. The level length is estimated by the median length over all the players. **d** For each level, the effect size of the environment on the normalized trajectory length as a function of the level length median. Positive Hedge’s g corresponds to longer trajectories (i.e. worse wayfinding performance) for city participants. **e** - Screenshots from City Hero Quest (CHQ), a city-themed version of SHQ. Map and image show 1 of the 5 levels tested. **f** - Association between Environment and trajectory lengths in SHQ and CHQ (N = 599). The center values correspond to the means. In all panels error bars correspond to 95% confiden6ce intervals.

People living in city centres typically travel shorter distances than people living in suburbs or in more rural environments, resulting from the denser arrangement of local activity locations [48, 49]. Thus, we hypothesized that city participants will have better wayfinding performance at SHQ levels requiring shorter trajectories. To test this hypothesis, we normalized the participants’ trajectory lengths at each level (*M* = 0, *SD* = 1) and plotted them against the corresponding game level length median, taken over all the participants. Figure 4c shows that the performance (inversely related to the normalized trajectory lengths) of city participants decreases with the game level length median, while it increases for non-city participants. Figure 4d shows a positive correlation between the effect size (Hedge’s g) of the environment on normalized trajectory length and the level length median (Pearson’s r = 0.50, *p* < 0.001). We computed a multiple linear regression with the environment effect size as the response variable, the level entropy and trajectory length median as the predictors. We found that both entropy (*F*_1,39_ = 5.20, *p* = 0.02, *η*^2^ = 0.12) and median trajectory length (*F*_1,39_ = 13.71, *p* < 0.001, *η*^2^ = 0.26) are significant predictors of the environment effect size.

### Replication of the association between environment and spatial ability in a city setting: City Hero Quest

While our virtual levels varied entropy from open waters to narrow inlets (see Supplementary Video 1), the SHQ navigation task was simulated in rural settings (rivers and ocean terrain) and may potentially favor participants who reported growing up outside cities. To test this hypothesis and directly replicate the environment effect with an independent sample, we designed a city-themed version of SHQ called City Hero Quest (CHQ) and tested participants with it alongside SHQ. In CHQ, the players performed the same task as in SHQ, but driving a car in city streets, see Figure 4e and Supplementary Video 2. We collected data from new participants on 5 SHQ levels and 5 CHQ levels matched for entropy and difficulty in order to test whether our SHQ results transfer to a city context, see Extended Data Fig. 7b-d. 599 participants were recruited in the United States via the crowdsourcing platform Prolific. We chose to collect data from the US as it was the most represented country in the Prolific participant pool, and a country with a high environment effect size in the initial SHQ dataset. Sample size justification and full description of this new task are available in the Supplementary Methods. The same data analysis pipeline was applied to Sea Hero Quest and City Hero Quest data. As shown in Figure 4f, the effect size of the environment on the normalized trajectory length is similar with SHQ data (Hedge’s g = 0.27, 95%CI=[0.06, 0.47]) and CHQ data (Hedge’s g = 0.34, 95%CI=[0.14, 0.54]), positive values indicating an advantage (i.e. shorter trajectories) for participants who grew up in non-city environments. The difference between the CHQ and SHQ environment effect was not significant (see Supplementary Information, CHQ Data Analysis section). This is also consistent with the effect size we found in the initial SHQ dataset when considering participants in the US on the same levels (Hedge’s g=0.30, 95%CI=[0.18, 0.42]). Because participants also provided their current environment (city/non-city), we were able to show that the effect of the current environment on CHQ or SHQ performance did not reach significance (see Supplementary Information). This suggests that the childhood period is key to predicting future spatial ability.

## Conclusions

Exploring population-level cognitive performance in 38 countries, we reveal that people are better at navigating in environments topologically similar to where they grew up. We show that this association is independent of age, gender, video games skill and education. Participants who grew up in less entropic cities show better performance at less entropic game levels, while participants who grew up in more entropic cities are better at navigating more complex game levels. Similarly, participants who grew up in cities generally perform better in game levels in smaller spaces than they do in game levels in larger spaces, while participants who grew up outside cities are better in larger game levels than in game levels in smaller spaces. These results support the idea that humans develop navigation strategies aligned with the type of environment they are exposed to, which become sub-optimal in other environments (see Supplementary Discussion). It indicates that the environment one grew up in is associated with cognitive ability, and that this association is stable across the life-span. Future research will need to explore how these differences emerge during childhood through adolescence, where abrupt changes in ability can occur [50].

## Methods

### Data

The design and the data collection process of Sea Hero Quest have been thoroughly described in [16].

#### Video game

In this study we focused on the wayfinding task. At the beginning of each wayfinding level, participants were shown locations (checkpoints) to visit on a map. The map disappeared, and they had to navigate a boat through a virtual environment to find the different checkpoints. Checkpoints were generally not encountered in the order of passage, but rather have to be navigated to by returning form one checkpoint to another (Figure 1). Participants were encouraged to collect as many ‘stars’ as possible across the levels: the faster the more stars were obtained. The first two levels were tutorial levels to familiarise the participant with the game commands.

#### Participants

A total of 3,881,449 participants played at least one level of the game. 60.8% of the participants provided basic demographics (age, gender, home country) and 27.6% provided more detailed demographics (home environment, level of education, see Methods). To provide a reliable estimate of spatial navigation ability, we examined the data only from participants who had completed a minimum of eleven levels of the game (including the first 4 wayfinding levels: levels 6, 7, 8 and 11) and who entered all their demographics. We removed participants above 70 years old because we previously showed a strong selection bias in this group causing their performance to be substantially higher than would be expected in unselected participants of the same age [16]. We also removed participants from countries with fewer than 500 players, or with education or environment levels more than 10-fold imbalanced. This resulted in 397,162 participants from 38 countries included in our analysis, (see Table S1).

#### Demographic information

Participants were made aware of the purpose of the game within the opening screen. Demographics were provided by consenting participants in two steps. First, their age, gender and home country were asked. Then, after having played a few levels, participants were invited to provide further information such as their level of education and the type of environment they grew up. They were asked whether they were willing to share their data with us and were guided to where they can opt out. The opt out was always available in the settings. Among the included participants there were 212,143 males (mean age: 37.81 ± 13.59 years old) and 185,173 females (mean age: 38.67 ± 14.92 years old). The levels (N = sample size) of education were: university (N=166,714), college (N=111,463), high-school (N=107,849), and no formal (N=11,290). We merged the university with the college levels due to their ambiguous meaning in some countries, and the high-school with the no formal level due to the relative low sample size of the latter. Hence, in our analysis the education variable had two levels: secondary and lower (N=119,139) and tertiary (N=278,177). The levels of home environment were: city (N=109,111), suburbs (N=131,738), mixed (N=80,266), rural (N=76,047). We merged the mixed, suburb and rural levels together to facilitate the interpretation of the effect of growing-up in city (N=109,111) and non-city (N=288,051) environments. City environments are distinguished from other settings due to the higher propensity for active, self-propelled travel (e.g. walking, cycling), relative to passive, car - or public transport-based travel [51], resulting from the denser arrangement of local activity locations. We furthermore anticipate that where participants have stated growing up in a city environment that there is a definitive and salient personal association with a city underlying this selection, whereas in other cases (e.g. suburban, mixed) this association is less clear, and better described as non-city. Both factors are common across international settings. Indeed, the observed clear difference between the city group and the other three groups, but little dissociation between the other three groups supports this approach, see Extended Data Fig. 2. To further validate this city / non-city dichotomy, we asked the participants to the City Hero Quest follow-up experiment the street they lived on in their home environment, and computed the entropy of the street network (SNE) in a spatial window around it (see Supplementary Methods). We then compared the SNE of people who reported having grown up in a city (N=114), suburb (N=326), rural (N=84) or mixed (N=75) environment. As shown in Extended Data Fig. 4c, the SNE in cities was significantly lower than the SNE in the other environments. We ran a one-way ANOVA with the reported home environment as independent variable and the SNE as dependent variable and found a significant effect of home environment (F(1,3)=25.72, p<0.001). Post-hoc pairwise t-test Bonferroni-corrected for multiple comparisons showed that the SNE of a city environment was significantly lower than the SNE of all the other environments (all p<0.001). We found no significant difference in the SNE of mixed vs rural (p=1), suburbs vs. mixed (p=0.20), and a small difference in the SNE of suburbs vs. rural (p=0.02).

For the analysis on the game level entropy, we included the participants that played all the Sea Hero Quest wayfinding levels (N = 10,626). There were 5,219 males (age: 41.89 *±* 15.95 years old), 5,407 females (age: 41.98 *±* 16.32 years old), 7,429 with tertiary education, and 3,604 grew up in cities.

#### Behavioural data

We collected the trajectory of each participant across each level. The coordinates of participants’ trajectories were sampled at Fs = 2 Hz.

### Metrics

#### Geospatial analysis

We focused on the quantification of the structural complexity of larger cities instead of the complexity of areas outside cities. This is because city streets can be more strictly compared with one another. On the opposite, areas outside cities can be heterogeneous both within and between countries, which makes the country-level averaging of their parameters problematic.

#### Street Network Entropy

We used the OSMnx toolbox to download the street network topology of cities from OpenStreetMap (OSM) [25]. For each included city we created a street network graph from OSM data within a 1000 × 1000 square meter box around the city geographical center. The use of a bounding box in the city centre is interesting as it is reflective of the wider city structure, and avoids issues related to classifications of regions, and administrative boundaries. This definition also has stronger persistence over time (considering city growth during the theoretical period of our analysis) [52]. To define the city geographical centers, we used the (latitude, longitude) coordinates provided by OpenStreetMaps. Then, we computed a 36-bin edge bearings distribution (1 bin every 10 degrees), taking one value per street segment. We initially took twice as many bins as desired, then merged them in pairs to prevent bin-edge effects around common values like 0 and 90 degrees. We also moved the last bin to the front, so e.g. 0.01 degree and 359.99 degrees were binned together. We calculated the Shannon entropy of the city’s orientations’ distribution.

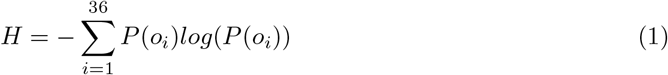

where i indexes the bins, and P(*o*_*i*_) represents the proportion of orientations that fall in the *i*^*th*^ bin [43]. For each of the 38 countries included in our analysis, we defined the average Street Network Entropy (SNE) as

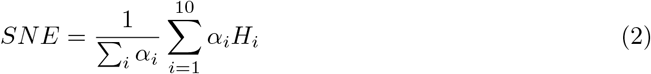

where (*H*_*i*_)_*i*∈[1..10]_ are the Shannon entropies of the 10 biggest cities in terms of population, and *α*_*i*_ is the population of the *i*^*th*^ city (see Table S1).

Since OSM mapping relies on the contributions from volunteers, we considered that this could introduce a bias, some countries being more densely mapped than others. So we compared these SNE values to the ones based on the city centers (latitude, longitude) coordinates provided by Google Maps (GM). We computed SNE value both from the drivable public streets network (‘drive’) and from the all non-private streets and paths network (‘all’). The ‘drive’ network is a more reliable and consistent source of long-term street network data, given that it represents the major established roads in each city. The ‘all’ network, by additionally covering pathways and pedestrian zones, is more susceptible to between-country variation in volunteer mapping practices and recent planning initiatives. We found little variations, see Table 1. We also varied the size of the street network box around the city centers. If the bounding box were too big it could go beyond the city boundaries (especially for smaller cities), but if too small it might not be representative of the whole city. We computed SNEs for 500 × 500, 1000 × 1000, 2000 × 2000 and 5000 × 5000 square meter boxes. Again, our results remained stable across the different sizes, see Table 1.

**Table 1.** Pearson’s correlation between environment effect size and different SNE calculations. The SNE set in bold is the one used in this manuscript. OSM = OpenStreetMaps, GM = Google Maps. P-values are from a t-test testing the hypothesis of no correlation against the alternative hypothesis of a nonzero correlation.

|  | City centre | Bounding box size (m) | Type of street network | correlation |
| --- | --- | --- | --- | --- |
| SNE1 | GM | 500 | drive | $r=-0.60, p < 0.001$ |
| SNE2 | GM | 500 | all | $r=-0.52, p = 0.001$ |
| **SNE3** | **OSM** | **1000** | **drive** | $r=-0.60, p = 0.001$ |
| SNE4 | OSM | 1000 | all | $r=-0.57, p = 0.001$ |
| SNE5 | GM | 1000 | drive | $r=-0.64, p = 0.001$ |
| SNE6 | GM | 1000 | all | $r=-0.58, p = 0.001$ |
| SNE7 | GM | 2000 | drive | $r=-0.58, p = 0.001$ |
| SNE8 | GM | 2000 | all | $r=-0.53, p = 0.001$ |
| SNE9 | GM | 5000 | drive | $r=-0.54, p < 0.001$ |
| SNE10 | GM | 5000 | all | $r=-0.50, p = 0.001$ |

#### Graph-theoretic metrics

Graph based measures are calculated on the ‘primal’ representation of the road network for each study area, where junctions are represented by nodes and roads as edges. This representation is typical in the calculation of road network-based graph metrics. The metrics selected were chosen on the basis of their widespread use in describing street networks, and full description of their implementation can be found in [25].

#### Circuity

This measures the of the sum length of all edges divided by the sum of Euclidean distances between nodes. Thus it captures the extent of deviation required from the most direct route when moving between two points on a network [53].

#### Mean Cluster Coefficient

This measure records the ratio of number of connections with neighbours over the total of number of possible connections, taken as a mean for all nodes. As such, it captures the degree close to which there is high interconnectivity between nodes in a network [54].

#### Mean Closeness Centrality

Closeness represents how close a node is to all other nodes within a network. It has been demonstrated to align closely with locations of activities [55] and correlated with activity in the anterior hippocampus when navigating [56]. The mean value for all nodes is taken.

#### Mean Betweenness Centrality

Betweenness centrality measures the extent to which a node features on shortest paths between all node pairs. Again, we use the mean value for the network, which indicates the extent to which flows of people may be spread or concentrated across the network [40].

#### Mean Degree Centrality

This measure records the fraction of nodes that each node is connected to, taken as a mean for all nodes in the network. This measure reflects the extent of connectivity between nodes on a network [57], is suitable for analysis interconnectivity in small areas and correlated with posterior hippocampal activity during navigation [56].

#### Mean Neighbor Degree

This measures for each node the mean degree of all neighboring nodes, and reflects the degree of local node connectivity.

#### Mean Street length

This measures the mean length of street segments, and thus is an indicator of block length. This provides a measure of the density of the street network.

#### Route-based metrics from agent-based simulations

All routes were calculated based on a ‘dual’ representation of the road network, whereby road segments are modelled as vertices and costs between vertices (e.g. distance, angular change) modelled as network edges. The Djikstra algorithm was used for identifying the optimal paths, with road length used as the optimisation measure. For each city, 1000 routes were generated for two randomly selected origin and destination nodes (i.e. road segment centre points). For each path, the following measures were extracted.

#### Unique streets

Sum of the unique street names provided by Open Street Map encountered along the route.

#### Partitions crossed

Sum of unique partitions encountered during the route. The road network was partitioned using the Louvain community detection algorithm on the dual graph, setting edge cost as angular change. These partitions have been used a proxy for deriving perceived neighborhood boundaries [58], and have demonstrated consistency with well-known regions, such as Soho and Mayfair in London, see Extended Data Fig. 10.

#### Snapped angular change

Angular deviations are calculated as the angle of incidence between two adjacent road segments, based on the connecting straight-line segments on each road polyline. The sum of absolute angular change between two consecutive road segments along a route from zero or 90 degrees (whichever is closer). We examined this novel measure because past work has shown that spatial memory for target locations is better after 90 degree or 180 rotations than other angular changes [59]. Memory for the angle of turns is biased towards right angles [60]. This suggests that it is easier to develop precise memories for low-SNE cities than for high-SNE cities. High-SNE cities would then require more training/learning, thus training navigation abilities.

#### Turns

Sum of turns between two consecutive segments surpassing 50 degrees in either direction, with more turns indicative of higher perceived cognitive distance [61]. We computed the same metric with 60 degrees, the results remained stable.

#### Angular deviation from target

The sum of the angular deviation from the destination, recorded at each road segment. Specifically this is recorded as the sum of differences between the angular change between two consecutive segments and the angular direction of the target from the first segment. During navigation angular deviation from the target positively correlates with activity in the posterior parietal lobe [62–64]. The posterior parietal lobe is a core part of the brain regions needed for effective navigation of familiar places [47].

### Video game analysis

#### Wayfinding performance

As in [16], we computed the trajectory length in pixels, defined as the sum of the Euclidean distance between the points of the trajectory. To control for familiarity with technology, we normalised the trajectory lengths by dividing them by the sum of their values at the first two levels. The first two levels only reflected video gaming skill (motor dexterity with the game controls) as no sense of direction was required to complete them. We defined an overall wayfinding performance metric corrected for video gaming skill (*W F*) as the 1st component of a Principal Component Analysis across the normalized trajectory lengths of the first four wayfinding levels (levels 6, 7, 8 and 11, 60.5% of variance explained). This metric being based on the trajectory length, it varies as the opposite of the performance: the longer the trajectory length, the worse the performance. We took the additive inverse of the metric and added an offset, so that *WF* = 0 corresponds to the worst performances. Pearson’s correlation coefficient between WF and the sum of the trajectory lengths of the first two levels (video gaming skills) is weak: *r* = 0.10, *p* < 0.001, bootstrapped 95%CI = [0.09, 0.10]. The implementation of WF is available in the code presented in the *Code and Data Availability* section.

#### Game Level Entropy

We calculated the Shannon entropy of the Sea Hero Quest level’s orientations’ distribution similarly as for the cities. To create the equivalent of “streets” in the levels of the game, we computed the Voronoi regions from the game level’s layout, and took their edges. The Voronoi region boundaries are considered equivalent to road centre lines in the city context. We then used the Douglas-Peucker algorithm to simplify the line made of the connected segments [65], see Figure 4a. For all game levels, we used a maximum offset tolerance between the original and the simplified line of three pixels. The entropy of the orientation distribution of the Game Level’s segments was then computed with equation 1.

### Statistical analysis

Further details are available in the Supplementary Information.

#### Linear Mixed Model computation

The parameters of the linear mixed models have been estimated with the maximum likelihood method (ML), and the covariance matrix of the random effects have been estimated with the Cholesky parameterization.

#### Low-SNE and High-SNE country clustering

We partitioned the 38 countries into two clusters based on their SNE with the k-means algorithm. We used the squared Euclidean distance metric. We ran the algorithm 1000 times and the arrangement never changed. The first group (low-SNE) comprised Australia, Canada, South Africa, Saudi Arabia, the United Arab Emirates, the United States of America and Argentina, with mean SNE = 2.69, SD = 0.18. The second group (High-SNE) comprised all the other countries, with mean SNE = 3.30, SD = 0.13.

#### Hedge’s g

Hedge’s g between group 1 and group 2 is defined as

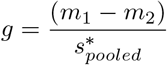

with *m*_*i*_ the mean of group *i*, and 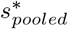 the pooled and weighted standard deviation:

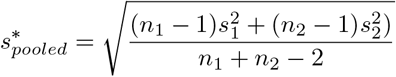

with *n*_*i*_ the sample size of group *i*, and *s*_*i*_ the standard deviation of group *i*.

The 95% confidence intervals displayed in this manuscript are exact analytical confidence intervals based on iterative determination of noncentrality parameters of noncentral t or F distributions. For more details, see [66].

#### Confidence Intervals (CI) for Pearson’s correlation coefficient

To estimate the uncertainty around Pearson’s correlation coefficient, we computed its percentile bootstrapped 95% CI. At each iteration, we resampled pairs of observations with replacement and computed their correlation values. We iterated this process 1000 times. We then sorted the correlation values and took the 2.5 and 97.5 percentiles obtained to yield a 95% CI. We illustrated this process for the correlation between Street Network Entropy and Environment effect size in Extended Data Fig. 4.

#### Linear regression predicting environment effect sizes (country slopes) based on SNE and GDP per capita

A multiple regression was calculated to predict the environment slopes based on SNE and GDP per capita. A significant equation was found (F(2,35)=22.40, *p* < 0.001) with a *R*^2^ = 0.56: environment slopes = 0.28 + 8.5 × 10^−7^(GDP) - 0.09(SNE). Both SNE and GDP per capita were significant predictors of environment slopes, see Table 2.

**Table 2.** Linear regression predicting environment slopes based on SNE and GDP per capita. Number of observations: 38, Error degrees of freedom: 35 Root Mean Squared Error: 0.0279 R-squared: 0.561, Adjusted R-Squared 0.536.

|  | Estimate | Standard Error | t-value | p-value |
| --- | --- | --- | --- | --- |
| (intercept) | 0.296 | 0.0528 | 5.61 | $2.53 \times 10^{-6}$ |
| SNE | -0.0976 | 0.0166 | -5.86 | $1.18 \times 10^{-6}$ |
| GDP | $9.76 \times 10^{-7}$ | $2.43 \times 10^{-7}$ | 4.01 | $2.96 \times 10^{-4}$ |

#### Modeling the environment effect with SNE vs all the other metrics

Two linear regressions were calculated to predict the environment slopes based on

1- SNE only (model ‘SNE only’): Env-slope ∼ SNE

We found a significant equation (F(1,36)=20.1, *p* < 0.001), Adjusted R-Squared = 0.341, BIC = -142.11.

2- All the other metrics (model ‘other metrics’): Env-slope ∼ unique-streets + turns + partition-crossed +dev-from-90-turns + dev-from-targets + street-length + neighbor-degree + circuity + clustering-coefficient + closeness-centrality + betweenness-centrality + degree-centrality.

We did not find a significant equation (F(10,25)=1.44, *p* = 0.21), adjusted R-Squared = 0.125, BIC = -105.18.

## Data Availability

Due to its considerable size (∼ 1 To), the global dataset is available on a dedicated server: https://shqdata.z6.web.core.windows.net/. A login/password to the server will be provided from the corresponding author upon request. A lighter version with the preprocessed trajectory lengths and demographic information is available at https://osf.io/7nqw6/?view_only=6af022f2a7064d4d8a7e586913a1f157

We also set up a portal where researchers can invite a targeted group of participants to play Sea Hero Quest and generate data about their spatial navigation capabilities. Those invited to play the game will be sent a unique participant key, generated by the Sea Hero Quest system according to the criteria and requirements of a specific project. https://seaheroquest.alzheimersresearchuk.org/

## Code Availability

The code allowing to reproduce the presented analyses is available along the the preprocessed trajectory lengths and demographic information at https://osf.io/7nqw6/?view_only=6af022f2a7064d4d8a7e586913a1f157.

## Ethics

This study has been approved by UCL Ethics Research Committee. The ethics project ID number is CPB/2013/015

## Acknowledgments

The authors wish to thank Deutsche Telekom for supporting and funding this research, Alzheimer Research UK (ARUK-DT2016-1) for funding the analysis, the Glitchers Limited for the design and game production, Saatchi and Saatchi London for project management and creative input.

## Author Contributions

HS, MH and AC supervised the project, HS, MH, AC, SG, CG, RD, JW, and CH designed research; AC, EM, GF, and DY analyzed data; AC, EM and HS wrote the paper.

## Supplementary Video legends

**Supplementary Video 1** | Examples of navigation in two Sea Hero Quest levels: level 27 (left) and level 58 (right).

**Supplementary Video 2** | Example of navigation in one City Hero Quest level.

## References

1. Kempermann, G., Kuhn, H. G. & Gage, F. H. More hippocampal neurons in adult mice living in an enriched environment. Nature 386, 493–495 (1997).

2. Hackman, D. A., Farah, M. J. & Meaney, M. J. Socioeconomic status and the brain: mechanistic insights from human and animal research. Nature Reviews Neuroscience 11, 651–659 (2010).

3. May, A. Experience-dependent structural plasticity in the adult human brain. Trends in Cognitive Sciences 15, 475–482 (2011).

4. Van Praag, H., Kempermann, G. & Gage, F. H. Neural consequences of environmental enrichment. Nature Reviews Neuroscience 1, 191–198 (2000).

5. Freund, J. et al. Emergence of individuality in genetically identical mice. Science 340, 756–759 (2013).

6. Clemenson, G. D., Deng, W. & Gage, F. H. Environmental enrichment and neurogenesis: from mice to humans. Current Opinion in Behavioral Sciences 4, 56–62 (2015).

7. Kardan, O. et al. Neighborhood greenspace and health in a large urban center. Scientific Reports 5, 1–14 (2015).

8. Dadvand, P. et al. Green spaces and cognitive development in primary schoolchildren. Proceedings of the National Academy of Sciences 112, 7937–7942 (2015).

9. Engemann, K. et al. Residential green space in childhood is associated with lower risk of psychiatric disorders from adolescence into adulthood. Proceedings of the National Academy of Sciences 116, 5188–5193 (2019).

10. Berman, M. G., Stier, A. J. & Akcelik, G. N. Environmental Neuroscience. American Psychologist 74, 1039–1052 (2019).

11. Bratman, G. N. et al. Nature and mental health: An ecosystem service perspective. Science advances 5, eaax0903 (2019).

12. Lederbogen, F. et al. City living and urban upbringing affect neural social stress processing in humans. Nature 474, 498–501 (2011).

13. Kühn, S. et al. In search of features that constitute an “enriched environment” in humans: Associations between geographical properties and brain structure. Scientific Reports 7, 1–8 (2017).

14. Carey, I. M. et al. Are noise and air pollution related to the incidence of dementia? A cohort study in London, England. BMJ Open 8, 1–11 (2018).

15. Stier, A. et al. Rethinking Depression in Cities: Evidence and Theory for Lower Rates in Larger Urban Areas. medRxiv. doi.org/10.1101/2020.08.20.20179036 (2020).

16. Coutrot, A. et al. Global Determinants of Navigation Ability. Current Biology 28, 2861–2866 (2018).

17. Malanchini, M. et al. Evidence for a unitary structure of spatial cognition beyond general intelligence. npj Science of Learning 5, 1–13 (2020).

18. Spiers, H. J. & Maguire, E. A. Thoughts, behaviour, and brain dynamics during navigation in the real world. Neuroimage 31, 1826–1840 (2006).

19. Maguire, E. A., Woollett, K. & Spiers, H. J. London taxi drivers and bus drivers: a structural MRI and neuropsychological analysis. Hippocampus 16, 1091–1101 (2006).

20. Xu, J. et al. Global urbanicity is associated with brain and behaviour in young people. Nature Human Behaviour, 1–15 (2021).

21. Coutrot, A. et al. Virtual navigation tested on a mobile app is predictive of real-world wayfinding navigation performance. PLoS ONE 14, 1–15 (2019).

22. Spiers, H. J., Coutrot, A. & Hornberger, M. Explaining World-Wide Variation in Navigation Ability from Millions of People: Citizen Science Project Sea Hero Quest. Topics in Cognitive Science (2021).

23. Sutherland, R. J. & Hamilton, D. A. Rodent spatial navigation: at the crossroads of cognition and movement. Neuroscience & Biobehavioral Reviews 28, 687–697 (2004).

24. Epstein, R. A., Patai, E. Z., Julian, J. B. & Spiers, H. J. The cognitive map in humans: spatial navigation and beyond. Nature neuroscience 20, 1504 (2017).

25. Boeing, G. OSMnx: New methods for acquiring, constructing, analyzing, and visualizing complex street networks. Computers, Environment and Urban Systems 65, 126–139 (2017).

26. Coughlan, G. et al. Toward personalized cognitive diagnostics of at-genetic-risk Alzheimer’s disease. Proceedings of the National Academy of Sciences 116, 9285–9292 (2019).

27. Klencklen, G., Després, O. & Dufour, A. What do we know about aging and spatial cognition? Reviews and perspectives. Ageing Research Reviews 11, 123–135 (2012).

28. Lester, A. W., Moffat, S. D., Wiener, J. M., Barnes, C. A. & Wolbers, T. The Aging Navigational System. Neuron 95, 1019–1035 (2017).

29. Nazareth, A., Huang, X., Voyer, D. & Newcombe, N. A meta-analysis of sex differences in human navigation skills. Psychonomic bulletin & review 26, 1503–1528 (2019).

30. Ritchie, S. J. & Tucker-Drob, E. M. How Much Does Education Improve Intelligence? A Meta-Analysis. Psychological Science 29, 1358–1369 (2018).

31. Ulrich, S., Grill, E. & Flanagin, V. L. Who gets lost and why: A representative cross-sectional survey on sociodemographic and vestibular determinants of wayfinding strategies. PLoS ONE 14, 1–16 (2019).

32. Fuchs, F. et al. Exposure to an enriched environment up to middle age allows preservation of spatial memory capabilities in old age. Behavioural Brain Research 299, 1–5 (2016).

33. Lynch, K. The Image of the City (The Technology Press Harvard University Press, Cambridge, USA, 1960).

34. Marshall, S. Streets and Patterns (Spon Press, Abingdon, UK, 2005).

35. Watts, A., Ferdous, F., Diaz Moore, K. & Burns, J. M. Neighborhood integration and connectivity predict cognitive performance and decline. Gerontology and Geriatric Medicine 1, 1–9 (2015).

36. Koohsari, M. J. et al. Cognitive Function of Elderly Persons in Japanese Neighborhoods: The Role of Street Layout. American Journal of Alzheimers Disease Other Dementias 34, 381–389 (2019).

37. Bongiorno, C. et al. Vector-based pedestrian navigation in cities. Nature Computational Science 1, 678–685 (2021).

38. Boeing, G. A multi-scale analysis of 27,000 urban street networks: Every US city, town, urbanized area, and Zillow neighborhood. Environment and Planning B: Urban Analytics and City Science, 1–18 (2018).

39. Shannon, C. E. A mathematical theory of communication. Bell System Technical Journal 27, 379–423 (1948).

40. Barthélemy, M. Spatial Networks. Physics Reports 499, 1–101 (2011).

41. Gudmundsson, A. & Mohajeri, N. Entropy and order in urban street networks. Scientific reports 3, 1–8 (2013).

42. Batty, M., Morphet, R., Masucci, P. & Stanilov, K. Entropy, complexity, and spatial information. Journal of geographical systems 16, 363–385 (2014).

43. Boeing, G. Urban Spatial Order: Street Network Orientation, Configuration, and Entropy. Applied Network Science 67, 1–20 (2019).

44. McNamee, D., Wolpert, D. & Lengyel, M. Efficient state-space modularization for planning: theory, behavioral and neural signatures. Advances in Neural Information Processing Systems (NIPS), 4511–4519 (2016).

45. Wiener, J. M., Schnee, A. & Mallot, H. A. Use and interaction of navigation strategies in regionalized environments. Journal of Environmental Psychology 24, 475–493 (2004).

46. Brunyé, T. T. et al. Strategies for selecting routes through real-world environments: Relative topography, initial route straightness, and cardinal direction. PloS ONE 10, e0124404 (2015).

47. Ekstrom, A. D., Spiers, H. J., Bohbot, V. D. & Rosenbaum, R. S. Human spatial navigation (Princeton University Press, 2018).

48. Salon, D. Heterogeneity in the relationship between the built environment and driving: Focus on neighborhood type and travel purpose. Research in Transportation Economics 52, 34–45 (2015).

49. Lenormand, M., Bassolas, A. & Ramasco, J. J. Systematic comparison of trip distribution laws and models. Journal of Transport Geography 51, 158–169 (2016).

50. Nazareth, A., Weisberg, S. M., Margulis, K. & Newcombe, N. S. Charting the development of cognitive mapping. Journal of Experimental Child Psychology 170, 86–106 (2018).

## References

51. Montello, D. R. A conceptual model of the cognitive processing of environmental distance information in International Conference on Spatial Information Theory (2009), 1–17.

52. Masucci, A. P., Arcaute, E., Hatna, E., Stanilov, K. & Batty, M. On the problem of boundaries and scaling for urban street networks. Journal of the Royal Society Interface 12 (2015).

53. Giacomin, D. J. & Levinson, D. M. Road network circuity in metropolitan areas. Environment and Planning B: Planning and Design 42, 1040–1053 (2015).

54. Jiang, B. & Claramunt, C. Topological analysis of urban street networks. Environment and Planning B: Planning and design 31, 151–162 (2004).

55. Porta, S. et al. Street centrality and densities of retail and services in Bologna, Italy. Environment and Planning B: Planning and design 36, 450–465 (2009).

56. Javadi, A.-H. et al. Hippocampal and prefrontal processing of network topology to simulate the future. Nature Communications 8, 1–11 (2017).

57. Jiang, B. & Claramunt, C. A structural approach to the model generalization of an urban street network. GeoInformatica 8, 157–171 (2004).

58. Filomena, G., Verstegen, J. A. & Manley, E. A computational approach to ‘The Image of the City’. Cities 89, 14–25 (2019).

59. Mou, W., McNamara, T. P., Valiquette, C. M. & Rump, B. Allocentric and egocentric updating of spatial memories. Journal of experimental psychology: Learning, Memory, and Cognition 30, 142 (2004).

60. Tversky, B. Distortions in memory for maps. Cognitive Psychology 13, 407–433 (1981).

61. Sadalla, E. K. & Magel, S. G. The perception of traversed distance. Environment and Behavior 12, 65–79 (1980).

62. Spiers, H. J. & Maguire, E. A. A navigational guidance system in the human brain. Hippocampus 17, 618–626 (2007).

63. Howard, L. R. et al. The hippocampus and entorhinal cortex encode the path and Euclidean distances to goals during navigation. Current Biology 24, 1331–1340 (2014).

64. Spiers, H. J. & Barry, C. Neural systems supporting navigation. Current Opinion in Behavioral Sciences 1, 47–55 (2015).

65. Douglas, D. H. & Peucker, T. K. Algorithms for the reduction of the number of points required to represent a digitized line or its caricature. Cartographica: the international journal for geographic information and geovisualization 10, 112–122 (1973).

66. Hentschke, H. & Stüttgen, M. C. Computation of measures of effect size for neuroscience data sets. European Journal of Neuroscience 34, 1887–1894 (2011).

