## Supplementary Information for "Entropy of city street networks linked to future spatial navigation ability"

### Supplementary Discussion

In this study, we showed that people are better at navigating in environments topologically similar to where they grew up. This effect was modulated by the design of the cities in each country. Growing up in cities with simpler, grid-like cities lead to better results at video game levels with a regular layout, while growing up outside cities or in cities with more complex, organic street networks lead to better results at more entropic video game levels. A plausible interpretation is that people become "optimized" to the environment where they grew up. They might develop navigation strategies tailored for a specific type of environment, which become sub-optimal in other environments. The reasons behind these country-level differences in city street networks are mostly historical. For instance, in South America, grid city design is characteristic of Hispanic American colonization [67,68], while disorganized street networks correspond to the typical organic street pattern of old European city cores [69]. The effect of the environment layout on spatial ability echoes recent studies showing that environmental barriers (e.g. streets) compartmentalize humans' cognitive map, disrupting the way spatial information is encoded in the entorhinal cortex [70, 71]. Our results also echo another recent study showing that participants from Padua (Italy) have better navigation abilities than participants from Utah (USA), suggesting that more complex environments facilitate better navigation abilities [72]. In order to test a sufficient sample of participants it was necessary to use a virtual world implemented via a mobile app. Because virtual navigation lacks the self-motion cues experienced in the real-world, one could argue that such a task might be more predictive of video games skill than real-world navigation. However, we have previously shown that navigation performance in Sea Hero Quest is predictive of navigating in the real world [73].

**Limitations**

**Design of the task -** For many details of the Sea Hero Quest task and data collection please see our previous publication [74]. To explore spatial navigation ability via an app for mobile phones and tablet devices we opted to explore performance in a wayfinding task requiring navigating to a set of check-points after memorisation of a map, see Supplementary Video 1. A wide range of existing approaches to testing navigation were rejected in the design process due to the challenge of making these succificiently engaging in a video game played on a mobile phone. For example, Morris-Water-Maze styled environments/tasks would likely not be sufficiently engaging due to the repeated use of the same panorama and game play would involve much of the time continually turning to reach the goal. By testing navigation over 75 levels in the game with varying conditions we aim to explore the impact of the structure of the environment on navigation ability. The choice of a boat navigating water was suggested by the game developers (Glitchers Ltd) and was supported by our team because it would: a) clearly demarcate where the participant can and cannot go in the environment (important for older participants), b) allow the challenge of acquiring the controls to feel more natural - for most people learning to steer a new boat can be a challenge which is reflected in the game. The wayfinding task in Sea Hero Quest belongs to the path planning category defined in Wiener et al.'s taxonomy of human wayfinding tasks [75]. We focused on this task because by varying properties of the levels we were able to examine a wide range of elaborate reasoning processes including interpretation of a map, planning a multi-step route, memory of the route, monitoring progress along the route and updating of route plan (detours), and transformation of bird's-eye perspective to an egocentric perspective needed for navigation. However the Sea Hero Quest wayfinding task does not cover all types of spatial abilities [75], and further work is needed to test whether our findings generalize to other spatial tasks.

**Path integration task -** Sea Hero Quest also features a path integration task in several levels (flare levels). These were included every 5th level in the game after level 4. In these levels, participants navigate along a river with bends to find a flare gun and then choose which three directions is the correct direction back to the starting point. In this study, we did not focus on path integration data because providing a reasonable number of trials (minimum 4 trials) requires participants to play 19 levels of the game. This makes the sample size in this task much lower, and substantially reduces the number of countries that can be examined (182,122 participants from 26 countries). Moreover, the Path Integration Performance being binary (correct /incorrect, see Supplementary Methods), it is less sensitive to between-subject differences than the continuous Wayfinding Performance. Nonetheless we do examine the Path Integration data to assess the overall pattern, see the Supplementary Methods and the Extended Data Fig. 6.

**City complexity quantification -** A critical aspect of our approach lies in the quantification of city "complexity". Urban pattern analysis is a vast interdisciplinary field of research, spanning from sociology [76] to architecture [77] and traffic forecast [78]. If a city's culture and history influence its topology [79], the latter in return structures the behaviour, cognition and mental health of its inhabitants [80–82]. Space Syntax studies developed a set of metrics to quantify the different dimensions of the interactions between societies and urban patterns [83]. In this study we used the Shannon entropy of the street network orientations: the higher the entropy the more organic the city, the lower the more grid-like. This choice was motivated by the strong theoretical connections between entropy and many measures of complexity [84, 85]. Its implementation was enabled by OSMnx, which grants access to the street network of any city in the world [86]. However, the difficulty to navigate a city cannot be entirely captured by its topological complexity. Other factors, such as the presence of salient landmarks (e.g. the Eiffel Tower in Paris) can significantly help situate oneself in space [87]. Studying the interaction between street networks complexity and landmark saliency would be a fascinating endeavor, but it requires building a database quantifying the saliency of city landmarks across the world.

**Video game complexity quantification -** We showed that participants who grew up in low-entropy cities were better at low-entropy Sea Hero Quest wayfinding levels than their compatriots who grew up in non-city areas. It was not true for participants who grew up in higher-entropy cities. We quantified game level's entropy in the same way as for the cities (see Methods), which only captures a facet of level's complexity. Further research will be needed to quantify the influence of visibility or landmark saliency on wayfinding performance in virtual environments (e.g. see [88]), and its interactions with the participant's home environment complexity.

**Potential cultural effect -** Participants from different countries could have different definitions for urban and rural environments. Future studies could disambiguate this by collecting more information on the participant's upbringing environment and objectively describe it as city or rural. We did this with our City Hero Quest experiment in the US (Extended Data Fig. 4c), but it could be replicated in other countries.

**Other potential confounding variables -** Other factors associated with growing up in cities could also be driving the differences in navigation. For instance, the associations between GPS use, transport mode (walking, driving, cycling, public transports...) environment and performance at our tasks would be worth exploring. Our analyses indicate that neither education or video games skill was a factor underlying the differences between growing up in a city or non-city. Future studies using birth cohorts would be a useful way to address the impact of other demographic factors that may be correlated with growing up in a city. In particular, the influence of the profession of the participants, likely different in city and non-city environments, would be an interesting variable to investigate.

### Supplementary Notes

**Who is driving the effect of environment?**

Understanding which group (city or non-city participants) drives the measured environment effect is important. This is a difficult task because the environment effect needs to be jointly modeled with the influence of age, gender, and education. Simply comparing the raw data performance could be misleading because environment classes (city / non-city) are not necessarily balanced in age or gender. For instance, non-city participants could show better raw wayfinding performance than non-city participants simply because they are younger. In this section, we propose two different approaches to overcome this issue.

First, we compared the amount of variance explained by the difference between countries in Wayfinding Performance when only considering city or non-city participants. If the city participants are the ones driving the effect, we might expect more between-country variance for this group than for the non-city participants. To test this hypothesis, we fitted two linear mixed models (LMM): one with the city participants ($\text{LMM}_{\text{city}}$, the other with the non-city participants ($\text{LMM}_{\text{noncity}}$). Both models had the same structure: age, gender and education as fixed effect, country as random effect.

$$WF \sim age + gender + education + (1|country)$$

For both models, we computed the Variance Partition Coefficient (VPC), measuring the amount of variance in Wayfinding Performance explained by the difference between countries, after correcting for fixed effects. If the topology of cities were the only factor differentiating countries, $\text{VPC}_{\text{city}}$ should be larger than $\text{VPC}_{non-city}$. We measured $\text{VPC}_{non-city} = 2.45\%$ 95%CI=[1.38% 3.46%] and $\text{VPC}_{\text{city}} = 2.96\%$ 95%CI=[1.81% 4.21%]. To compute the confidence intervals around the VPC, we created a bootstrapped distribution of the VPC (500 iterations), then obtained the relevant quantiles from that distribution, see Extended Data Fig. 5a. We compared the bootstrapped distributions of $\text{VPC}_{\text{city}}$ and $\text{VPC}_{\text{noncity}}$ with an unpaired Student's t-test: t(998)=12.17, $p < 0.001$.

Second, we explored the link between the game level entropy and the difference in performance between city and non-city participants from low-SNE and high-SNE countries. If the environment effect was driven by the city structure, then city participants from low-SNE countries would be better at lower entropy game levels, while city participants from high-SNE countries would be better at higher entropy game levels. Conversely, this association should be absent in non-city participants. To test this hypothesis, we fitted four LMM: one with the city participants from low-SNE country ($\text{LMM}_{\text{city\_lowSNE}}$, one with the non-city participants from low-SNE country ($\text{LMM}_{\text{noncity\_lowSNE}}$), one with the city participants from high-SNE country ($\text{LMM}_{\text{city\_highSNE}}$), and the last with the non-city participants from high-SNE country ($\text{LMM}_{\text{noncity\_highSNE}}$). They all had the same structure with age, gender and education as fixed effect, and level number as random effect:

$$WF \sim age + gender + education + (1|level\_number)$$

We then computed the difference between the level conditional modes of $\text{LMM}_{\text{city\_highSNE}}$ and $\text{LMM}_{\text{city\_lowSNE}}$ on one side, and of $\text{LMM}_{\text{noncity\_highSNE}}$ and $\text{LMM}_{\text{noncity\_lowSNE}}$ on the other side. Let

$D_{\text{city}}$ = conditional_mode(level | $\text{LMM}_{\text{city\_highSNE}}$) - conditional_mode(level | $\text{LMM}_{\text{city\_lowSNE}}$)
and
$D_{\text{noncity}}$ = conditional_mode(level | $\text{LMM}_{\text{noncity\_highSNE}}$) - conditional_mode(level | $\text{LMM}_{\text{noncity\_lowSNE}}$)
For each level, $D_{\text{city}}$ represents the difference in Wayfinding Performance between city participants from high-SNE countries and city participants from low-SNE countries. For each level, $D_{\text{noncity}}$ represents the difference in Wayfinding Performance between non-city participants from high-SNE countries and non-city participants from low-SNE countries. Extended Data Fig. 5 panel b shows the relation between $D_{\text{city}}$ and level entropy, while Extended Data Fig. 5 panel c shows the relation between $D_{\text{noncity}}$ and level entropy.
The difference between high-SNE and low-SNE country effect was not significantly correlated with level entropy in non-city participants (Pearson's correlation, $r_{\text{noncity}} = -0.33, p = 0.053$). In city participants it was positively correlated ($r_{\text{city}} = 0.44, p = 0.009$). The correlation slope difference was significant (Fisher's z = 3.60, $p < 0.001$, 95% CI for $r_{\text{city}} - r_{\text{noncity}} = [0.36, 1.10]$). $D_{\text{city}}$ values were negative for low entropy game levels, meaning that growing up in low-SNE cities leads to better wayfinding ability in these levels. As game level entropy increases, $D_{\text{city}}$ becomes positive, meaning that city participants from high-SNE countries are better at more entropic game levels.
We also tested whether the correlations between environment effect size and level entropy (Figure 4b) was driven by a small number of anglophone countries such as the US, Australia, Canada, and the UK, which account for a large fraction of the total dataset. With the US, Australia and Canada in the low-SNE group, the correlation coefficient between the entropy of the levels and the low-SNE environment slopes was $\text{r}_{\text{lowSNE}} = 0.57$, p < 0.001. Between the entropy of the levels and the high-SNE environment slopes it was $\text{r}_{\text{highSNE}} = 0.44$, p = 0.003. Now, without the US, Australia and Canada in the low-SNE group (i.e. only keeping Argentina, Saudi Arabia, South Africa and the United Arab Emirates) we have $\text{r}_{\text{lowSNE}} = 0.55$, p < 0.001. This shows that this effect is not driven by a small number of over-represented countries.
These results support the hypothesis that participants perform better in environments with a topology similar to their home environment. These results also evidence variation between countries for non-city upbringing participants. More work will be needed to uncover the variables behind the variations in performance in non-city environments.

**Interaction between the effects of age and environment**
As shown in Extended Data Fig. 1, the effect size of the environment remains stable across age, with a maximum amplitude of Hedge's g variation across age of 0.04. Estimating the statistical significance of the interaction between age and environment is difficult. Because of the magnitude of the dataset, most effects are likely to always be 'significant below the 0.001 threshold'. This is why we chose to focus on effect sizes, which are independent of sample size. Another way to assess the magnitude of the interaction between age and environment is to compare the proportion of variance explained by age in two independent multiple linear regression analyses (LR): one with the city participants ($\text{LR}_{\text{city}}$, N=109,111), the other with the non-city participants ($\text{LR}_{non\_city}$, N=288,051).
In $\text{LR}_{\text{city}}$, age has the strongest effect ($F_{1,109107} = 15579$, $p < 0.001$, $\eta^2 = 0.12$, Hedge's $g = 0.99$, 95% CI = [0.96, 1.00]), followed by gender ($F_{1,109107} = 5952.6$, $p < 0.001$, $\eta^2 = 0.05$, Hedge's $g = 0.46$, 95% CI = [0.44, 0.47]), and education ($F_{1,109107} = 260.78$, $p < 0.001$, $\eta^2 = 0.002$, Hedge's $g = 0.12$, 95% CI = [0.10, 0.13])). The Hedge's g of age is computed between participants under 25 years old (N=24,674) and above 55 years old (N=15,090).
In $\text{LR}_{\text{noncity}}$, age has also the strongest effect ($F_{1,288047} = 46354$, $p < 0.001$, $\eta^2 = 0.13$, Hedge's $g = 0.99$, 95% CI = [0.98, 1.01]), followed by gender ($F_{1,288047} = 14787$, $p < 0.001$, $\eta^2 = 0.04$, Hedge's $g = 0.44$, 95% CI = [0.43, 0.45]), and education ($F_{1,288047} = 433.4$, $p < 0.001$, $\eta^2 = 0.005$,

Hedge's $g = 0.14$, 95% CI = [0.13, 0.15])). The Hedge's g of age is computed between participants under 25 years old (N=63,361) and above 55 years old (N=44,888).
The effect sizes ($\eta^2$ and Hedge's g) of age in $LR_{city}$ and $LR_{noncity}$ are very close, with largely overlapping 95% CI for Hedge's g. The closeness between these effect sizes is consistent with the idea that the difference between the effect of age on spatial ability in city and non-city participants is not meaningful.

**Robustness of the environment effect to the rate of city participants**
The city and non-city classes are imbalanced, with 72.5% of the participants having grown up outside cities (87% in the US). However, controlling for the non-city rate did not impact the significance of the effect of the Street Network Entropy (SNE) on the environment effect size. We ran a multiple linear regression with the environment effect size as the dependent variable, the SNE and the non-city rate as the independent variables and found a significant effect of both the SNE (t = -5.79, $p < 0.001$) and the non-city rate (t = 5.18, $p < 0.001$). We didn't find a significant correlation between the non-city rate and the SNE (r = -0.02, p = 0.91).
To further establish that the imbalance between the city and non-city class did not bias our results, we randomly subsampled the non-city class to match the sample size of the city class in each country, and it did not modify the country-level effect sizes. For instance, in Romania, the original effect size (without non-city subsampling) was $g_{\text{original}} = -0.033$, with positive values indicating an advantage for non-city participants. The average over 1000 random subsampling of the non-city class was $g_{\text{balanced}} = -0.034$. Over all countries, we obtained a Pearson's correlation between $g_{\text{original}}$ and $g_{\text{balanced}}$ of r = 0.995, $p < 0.001$.

### Supplementary Methods

**Path Integration analysis**
**Task -** Sea Hero Quest also features a path integration task in several levels (flare levels). These were included every 5th level in the game after level 4. In these levels, participants navigate along a river with bends to find a flare gun and then choose which three directions is the correct direction back to the starting point.
**Participants -** We applied the same inclusion criteria as for the Wayfinding Performance (see Methods). We examined the data only from participants who had completed at least the first 4 path integration levels: levels 4, 9, 14 and 19). This resulted in 181,122 participants (84,639 females) from 26 countries.
**Path Integration Performance -** For each Path Integration level, the performance measure is binary (correct /incorrect). In order to keep the overall measure binary and to use straightforward logistic models, we defined the overall Path Integration Performance (PI) as follows: PI = 1 if the response to all four levels is correct (67,977 participants), PI = 0 otherwise (113,145 participants).
**Statistical Analysis -** We performed the same analysis as for the Wayfinding Performance, and it yielded similar results. Since the response variable is binary, we computed a multivariate logistic regression was calculated to predict Path Integration Performance based on age, gender, education and environment. Age has the strongest effect ($F_{1,181117} = 11259$, $p < 0.001$, $OR = 0.96$, Hedge's $g = 0.71$, 95% CI = [0.69, 0.72]), followed by gender ($F_{1,181117} = 8031.70$, $p < 0.001$, $OR = 2.57$ (baseline = female), Hedge's $g = 0.41$, 95% CI = [0.40, 0.42]), environment ($F_{1,181117} = 914.98$, $p < 0.001$, $OR = 0.70$ (baseline = non-city), Hedge's $g = 0.14$, 95% CI = [0.13, 0.15]), and education ($F_{1,181117} = 482.30$, $p < 0.001$, $OR = 0.78$ (baseline = tertiary), Hedge's $g = 0.10$, 95% CI = [0.09, 0.11]). The Hedge's g of age is computed between participants under 25 years old (N=41,446) and above 55 years old (N=27,404). Extended Data Fig. 6, panel a presents the effect of the environment on Path Integration Performance stratified by age and gender.
To quantify how Path Integration Performance and environment are associated across countries,

we fit a mixed-effects logistic regression (GLMM) for Path Integration Performance, with fixed effects for age, gender and education, and a random effect for country, with random slopes for environment clustered by country. Extended Data Fig. 6, panel b presents the environment slopes for each country, positive values indicate an advantage for participants raised outside cities. In terms of Hedge's g, this spectrum ranges from Slovakia (g = 0.04, 95%CI=[-0.02, 0.11])) to the United States (g = 0.22, 95%CI=[0.19, 0.25]), with positive values indicating an advantage for participants raised outside cities.

**Association between Path Integration Performance and Street Network Entropy -** As in Figure 2 panel c for Wayfinding Performance, Extended Data Fig. 6 panel c shows for each country the Street Network Entropy as a function of the Environment Effect Size (random slopes) on Path Integration Performance. Similar to our Wayfinding Performance result, we find that the Pearson's correlation between Street Network Entropy and Environment Effect Size is negative. However, it does not reach significance (r(24)=-0.40, 95%CI = [-0.62, 0.13], p = 0.05).

**Rationale for statistical analysis**

A distinctive feature of this work is the magnitude of the dataset, which is unusual in human behavior modelling studies. As stated in the main manuscript, the magnitude of the dataset makes most effects likely to be 'significant below the 0.001 threshold', even when their effect sizes are too small to be relevant. This is why for each model we computed, we reported classic statistics (F-values, p-values) along with effect sizes, that are independent of the sample size (Hedge's g, Pearson's correlation coefficient or random effect slopes). For each effect size we computed 95% confidence intervals or standard errors. The basis of our statistical analysis are linear mixed models as they have the great advantage to control for fixed effects while estimating random effects on a partition of the dataset (e.g. country-level clusters). Here, we used fixed effects to control for known sources of variance in the response variable (gender, age and education). We used random effects to directly compare the effect size of our key variable (environment) across different levels, which we hypothesized to have a clustering effect on the response variable. We used two clustering variables: countries and game levels, respectively in the models

$$WF \sim age + gender + education + (1 + environment \mid country) \tag{1}$$

and

$$WF \sim age + gender + education + (1 + environment \mid level) \tag{2}$$

We chose models with the simplest structure able to answer our research questions. To limit the risk of type II errors we only included variables for which we had strong a priori hypotheses coming from previous studies. We then used the environment effect size (the random slopes) associated with each clustering variable to compute further analysis at the level of the country or of the game level (e.g. Figure 2c and Figure 4b).

**City Hero Quest**

- *Overview* - Participants were recruited in the United States via the crowdsourcing platform Prolific. We chose to collect data from the US as it was the most represented country in the Prolific participant pool, and was one of the countries with the highest environment effect size in our initial SHQ dataset. After providing their informed consent and an initial screening (more details below), participants were redirected to a Gorilla webpage. Gorilla is a cloud based research platform supporting behavioral experiments online. Participants completed a demographic questionnaire, and were given instructions to first install Sea Hero Quest on their mobile, and then install City Hero Quest (CHQ) on their personal computer. Participants played 5 Sea Hero Quest levels: training level 1, and wayfinding levels 11, 32, 42, 68. Then, they played 5 City

Hero Quest levels, each modeled from a corresponding Sea Hero Quest map (level 1, 11, 52, 56, 67), see Extended Data Fig. 7b-d. Levels 1 and 11 were common to both CHQ and SHQ, they were chosen to accustom the participant with the task. The other levels were identical in layout, matching entropy and difficulty. After completing both tasks, participants emailed us their data along with their unique identifying number.

- *Description of City Hero Quest -* City Hero Quest is a desktop virtual reality game designed to parallel Sea Hero Quest within an urban setting. The game was built in Unity 3D physics games engine (https://unity.com/) with all virtual structures modeled in Blender 3D modeling software (`www.blender.org`). To control the back end and log experimental variables, we used Unity Experimental Framework (UXF), an open source set of packages for Unity 3D designed for human behavioral research. The game consists of five levels: one training level and four wayfinding levels. Each level in City Hero Quest was modeled from a corresponding Sea Hero Quest map. Prior to building each structural model for City Hero Quest, a model city environment was built with urban panels designed to wrap around any 3D polygon. To build the model for each level, we extracted the base map for the corresponding Sea Hero Quest level (1, 16, 52, 56, or 67) and constructed 3D polygons similar in height to the sea environment. After constructing these polygons for each isolated structure within the base map, we tessellated the urban panels on each equally split face of each polygon. These panels were subsequently textured using urban textures (e.g. brick, marble) sourced from Poliigon (`www.poliigon.com`) and baked for scene lighting to reduce the runtime demand on player platforms. This process was repeated for all levels. Once built and baked, each model was imported to Unity 3D within separate scenes corresponding to each level and scaled up by 50 units. For the player, we imported a basic Unity car model (pre-fab), as this model contained the features needed for the game baked in. We edited the features of the car to mimic that of the boat in Sea Hero Quest, i.e. turn right, turn left, and accelerate, and de-activated additional features, i.e. lights, engine sounds. For each level, the car [player] was set at an initial position and pointing direction similar to that of the corresponding Sea Hero Quest level such that, at runtime, all players viewed the same perspective of the environment. Player position and rotation variables were logged at each frame (0.036 seconds) using the UXF position tracking package. This data was subsequently saved to each player's computer, split according to level.

- *Participants -* Participants were recruited on Prolific (`www.prolific.co`) over August and September of 2021 with data collection split into eight batches. Each batch was prescreened using the same set of restrictions: United States as 'current country of residence', United States as 'Nationality', ethnicity provided by the user (not screened for any specified ethnicities), Prolific approval rate of greater than or equal to 90%, and Mac OS / Ubuntu / or Windows as 'Computer Operating System'. Finally, participants from previous batches were excluded from all subsequent batches. Prolific approval was determined by player upload of both sets of data – Sea Hero Quest and City Hero Quest – to an anonymous OneDrive server. Partial datasets were approved but excluded from analysis. All approved participants were paid at a standard rate of £7.50 per hour. To determine the sample size, we computed a power analysis based on the effect size of the home environment obtained in the initial SHQ dataset. Since we planned to test participants in the US, we used an effect size Hedge's $g = 0.20$ (in the initial dataset, $g = 0.19$, 95%CI=[0.17, 0.21]). We also took into account the imbalance between city and non-city participants. In the initial dataset in the US we had 12,931 city and 86,254 non-city participants, so a ratio of 0.13. Here we decided to use a ratio of 0.20 to increase the statistical power. The required sample size to detect an effect size of $g = 0.20$ with 2 imbalanced groups (20% - 80%) with 80% statistical power is N1=200, N2=800 (pwr R package, based on [89]). Because testing 1000 participants on

Prolific is time consuming due to the screening and technical assistance required, we performed a sequential analysis while the data collection was in progress. Sequential analyses make it possible to perform high-powered informative experiments while sparing human and economic resources. They are widely used in large-scale medical trials. At an interim analysis, data collection can be stopped whenever the results are convincing enough to conclude that an effect is present, while controlling the Type 1 error rate, see [90]. We used Pocock boundary [91], which lowers the alpha level to the same value for each interim analysis such that the overall alpha level remains .05. Here, we planned to perform 5 interim analyses, so 1 every 200 participants. This means that for the association between the home environment and spatial ability to be considered significant and to decide to stop the data collection, the alpha needs to be <0.1 for the first interim analysis, <0.2 for the second, <0.3 for the third, <0.4 for the fourth, or <0.5 for the complete dataset. A significant alpha threshold was met at the third interim analysis, both for CHQ ($p < 0.001$) and SHQ ($p = 0.016$). At this point, we collected complete data from 599 participants with a city/non-city ratio of 0.19 (114 city dwellers). There were 299 males and 300 females, the mean age was 27.0 years (SD=8.1 years). City dwellers grew up in 65 different cities across the US.

- *Data Analysis* - The same data analysis pipeline was applied to Sea Hero Quest and City Hero Quest data. As for the initial dataset, we normalized the trajectory length of each wayfinding level by the length of the training level (level 1) to account for gaming skills. These values were z-scored for each level to allow a direct comparison between levels of different lengths. To get an overall performance value for each participant, we defined the 'Normalized Trajectory Length' as these z-scored values averaged over the 4 wayfinding levels. Note that the Normalized Trajectory Length is inversely related to performance (the higher the worse). The Normalized Trajectory Lengths for SHQ and CHQ were significantly correlated ($r = 0.44$, $p<0.001$).
Two multivariate linear regressions (one for CHQ, one for SHQ) were calculated to predict Normalized Trajectory Length based on age, gender, education and upbringing environment. For SHQ, we found a significant effect of Age ($F(1,594)=40.78$, $p < 0.001$, Hedge's $g = 0.93$ 95%CI = [0.04 1.81]), Gender ($F(1, 594)=24.10$, $p < 0.001$, Hedge's $g = 0.31$ 95%CI = [0.15 0.47]), and Upbringing Environment ($F(1, 594)=6.41$ , $p=0.012$, Hedge's $g = 0.27$ 95%CI = [0.06 0.47]). Education did not reach significance ($F(1, 594)=1.58$, $p=0.21$). For CHQ, we found a significant effect of Age ($F(1, 594)=18.44$, $p < 0.001$, Hedge's $g = 1.44$ 95%CI = [0.55 2.33]), Gender ($F(1,594)=34.66$, $p < 0.001$, Hedge's $g = 0.39$ 95%CI = [0.23 0.55]), and Upbringing Environment ($F(1, 594)=13.83$, $p < 0.001$, Hedge's $g = 0.34$, 95%CI=[0.14, 0.54]). Education did not reach significance ($F(1, 594)=0.12$ , $p=0.73$).
To directly compare difference between the CHQ and SHQ environment effect size, we computed a linear mixed model with Age, Gender, Education, Upbringing Environment and Task (+interactions) as fixed effects and Participant ID as random effect (to take into account the fact that Task is a within-subject variable).

$$NormTrajLength \sim (Age + Gender + Education + Environment) * Task + (1|ParticipantID)$$

We found a significant effect of Age ($F(1, 594)=41.12$, $p < 0.001$), Gender ($F(1,594)=42.14$, $p < 0.001$, and Upbringing Environment ($F(1, 594)=14.14$, $p < 0.001$. Education did not reach significance ($F(1, 594)=0.92$ , $p=0.34$), nor did Task ($F(1, 594)=4.00$ , $p=0.05$). None of the interaction between Task and the other fixed effects reached significance (Task:Age $F(1, 594)=3.37$ , $p=0.07$; Task:Gender $F(1, 594)=0.87$ , $p=0.35$; Task:Education $F(1, 594)=0.66$ , $p=0.41$; Task:Environment $F(1, 594)=1.21$ , $p=0.27$). Thus, we found no association between the task design (CHQ or SHQ) and the effect of the environment on spatial ability.
We ran the same regressions, just replacing the Upbringing Environment with the Current Environment. For SHQ, we found a significant effect of Age ($F(1,594)=40.72$, $p < 0.001$, Hedge's $g =$

0.93 95%CI = [0.04 1.81]), and Gender (F(1, 594)=22.65, p $<$ 0.001, Hedge's g = 0.31 95%CI = [0.15 0.47]). Current Environment did not reach significance (F(1, 594)=2.51 , p=0.08, Hedge's g = 0.11 95%CI = [-0.06 0.29]), nor did Education (F(1, 594)=1.91, p=0.17). For CHQ, we found a significant effect of Age (F(1, 594)=19.13, p $<$ 0.001, Hedge's g = 1.44 95%CI = [0.55 2.33]), Gender (F(1,594)=30.05, p $<$ 0.001, Hedge's g = 0.39 95%CI = [0.23 0.55]). Current Environment did not reach significance (F(1, 594)=2.53, p = 0.08, Hedge's g = 0.19, 95%CI=[0.01, 0.37]), nor did Education (F(1, 594)=0.27 , p=0.61).

- *Entropy of city vs non-city environments* - We computed the Street Network Entropy (SNE) in a window around the home environment addresses given by the participants, in the same way as we did for the city centers in the initial dataset. We then compared the SNE of people who reported having grown up in a city (N=114), suburb (N=326), rural (N=84) or mixed (N=75) environment. We varied the size of the window depending on the street density, as in some rural areas there only are a couple of roads in a 1000 $\times$ 1000 square meter box around the participant's address, which biases the SNE computation. The size of the window was chosen to match the average number of street segments in all 4 reported home environments. For reported city environments we chose a 1000 $\times$ 1000 square meter box, as in the initial SNE analysis (mean number of segments = 592 (SD=247), for suburb environments we chose a 1200 $\times$ 1200 square meter box (mean number of segments = 567 (SD=305), for mixed environments we chose a 1250 $\times$ 1250 square meter box (mean number of segments = 583 (SD=448), and for rural environments we chose a 2000 $\times$ 2000 square meter box (mean number of segments = 583 (SD=698). We ran a one-way ANOVA with the reported home environment as independent variable and the SNE as dependent variable and found a significant effect of home environment (F(1,3)=25.72, $p < 0.001$). Post-hoc pairwise t-test Bonferroni-corrected for multiple comparisons showed that the SNE of the city environment was significantly lower than the SNE of all the other environments (all $p < 0.001$), see Extended Figure 4c. We found no significant difference in the SNE of mixed vs rural (p=1), suburbs vs. mixed (p=0.20), and a small difference in the SNE of suburbs vs. rural (p=0.02). For completeness we also tested the correlation between the raw SNE values computed around participants' home addresses and Normalized Trajectory Length and didn't find a significant relationship (for SHQ, r = -0.01, p = 0.80; for CHQ, r = -0.04, p = 0.31).

### References


67. Stanislawski, D. The origin and spread of the grid-pattern town. *Geographical Review* **36**, 105–120 (1946).

68. Outtes, J. Disciplining society through the city: the genesis of city planning in brazil and argentina (1894–1945). *Bulletin of Latin American Research* **22**, 137–164 (2003).

69. Kostof, S. *The city shaped: urban patterns and meanings through history* (Thames and Hudson, London, United Kingdom, 1991).

70. Horner, A. J., Bisby, J. A., Wang, A., Bogus, K. & Burgess, N. The role of spatial boundaries in shaping long-term event representations. *Cognition* **154**, 151–164 (2016).

71. He, Q. & Brown, T. I. Environmental barriers disrupt grid-like representations in humans during navigation. *Current Biology* **29**, 2718–2722 (2019).

72. Barhorst-Cates, E. M., Meneghetti, C., Zhao, Y., Pazzaglia, F. & Creem-Regehr, S. H. Effects of home environment structure on navigation preference and performance: A

comparison in veneto, italy and utah, usa. *Journal of Environmental Psychology* **74**, 101580 (2021).

73. Coutrot, A. *et al.* Virtual navigation tested on a mobile app is predictive of real-world wayfinding navigation performance. *PLoS ONE* **14**, 1–15 (2019).
74. Coutrot, A. *et al.* Global Determinants of Navigation Ability. *Current Biology* **28**, 2861–2866 (2018).
75. Wiener, J. M., Büchner, S. J. & Hölscher, C. Taxonomy of human wayfinding tasks: A knowledge-based approach. *Spatial Cognition & Computation* **9**, 152–165 (2009).
76. Roy, A. Urban informality: toward an epistemology of planning. *Journal of the american planning association* **71**, 147–158 (2005).
77. Katz, P., Scully, V. J. & Bressi, T. W. *The new urbanism: Toward an architecture of community* (McGraw-Hill, New York, USA, 1994).
78. Black, J. *Urban transport planning: Theory and practice* (Routledge, London, United Kingdom, 2018).
79. Smith, M. E. Form and meaning in the earliest cities: a new approach to ancient urban planning. *Journal of Planning History* **6**, 3–47 (2007).
80. Kühn, S. *et al.* In search of features that constitute an "enriched environment" in humans: Associations between geographical properties and brain structure. *Scientific Reports* **7**, 1–8 (2017).
81. Carey, I. M. *et al.* Are noise and air pollution related to the incidence of dementia? A cohort study in London, England. *BMJ Open* **8**, 1–11 (2018).
82. Engemann, K. *et al.* Residential green space in childhood is associated with lower risk of psychiatric disorders from adolescence into adulthood. *Proceedings of the National Academy of Sciences* **116**, 5188–5193 (2019).
83. Hillier, B. *Space is the machine: a configurational theory of architecture* (Space Syntax, London, United Kingdom, 2007).
84. Batty, M., Morphet, R., Masucci, P. & Stanilov, K. Entropy, complexity, and spatial information. *Journal of geographical systems* **16**, 363–385 (2014).
85. Boeing, G. Measuring the Complexity of Urban Form and Design. *Urban Design International* 1–21 (2018).
86. Boeing, G. OSMnx: New methods for acquiring, constructing, analyzing, and visualizing complex street networks. *Computers, Environment and Urban Systems* **65**, 126–139 (2017).
87. Ewing, R. & Handy, S. Measuring the unmeasurable: Urban design qualities related to walkability. *Journal of Urban design* **14**, 65–84 (2009).
88. Yesiltepe, D. *et al.* Computer models of saliency alone fail to predict subjective visual attention to landmarks during observed navigation. *Spatial Cognition & Computation* 1–28 (2020).
89. Cohen, J. *Statistical power analysis for the behavioral sciences* (Lawrence Erlbaum Associates, 1988).

90. Lakens, D. Performing high-powered studies efficiently with sequential analyses. *European Journal of Social Psychology* **44**, 701–710 (2014).

91. Pocock, S. J. Group sequential methods in the design and analysis of clinical trials. *Biometrika* **64**, 191–199 (1977).
