## Extended Data Figures for "Entropy of city street networks linked to future spatial navigation ability"

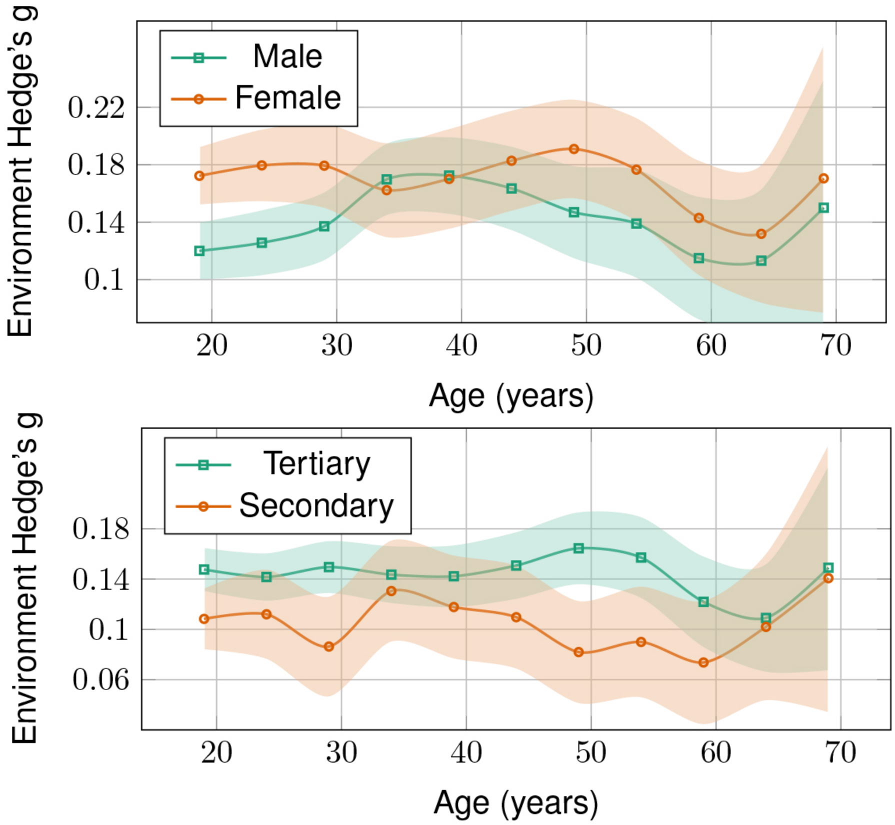


**Extended Data Figure 1. | Environment effect size across age, gender (top) and level of education (bottom).** Effect size is quantified with Hedge's g, within 5-year windows. Positive values correspond to an advantage for participants who grew-up outside cities. Error bars correspond to 95% confidence intervals and the center values correspond to the means.

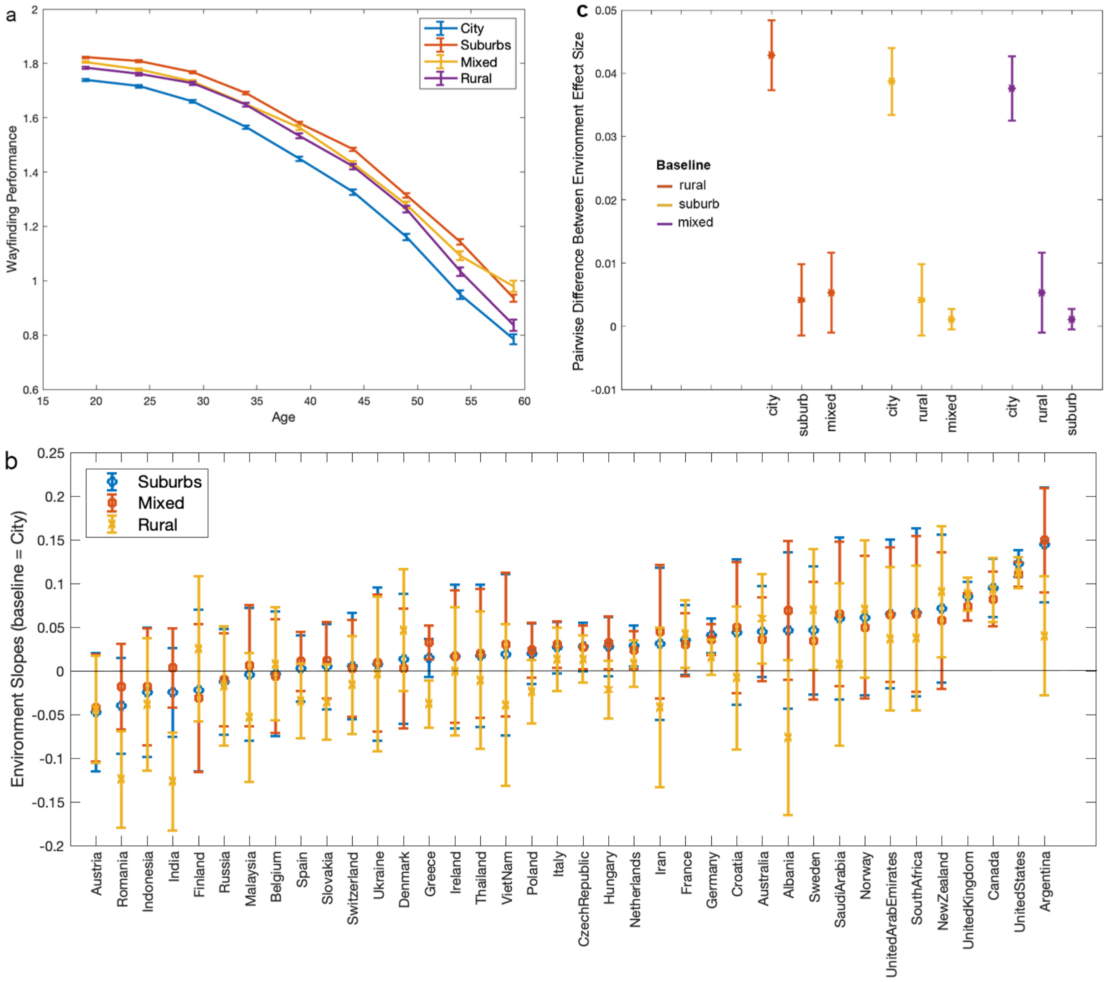


**Extended Data Figure 2.** | **Association between age, home environment, country, and Wayfinding Performance. a** - Wayfinding Performance as a function of age for participants who grew up in city, suburb, mixed and rural environments. Data points correspond to the wayfinding performance averaged within 5-year windows. **b** - Difference in the effect of growing up outside cities on wayfinding performance across countries. We fit a linear mixed model for wayfinding performance, with fixed effects for age, gender and education, and random environment slopes clustered by country, as in Figure 2a. Suburbs, Mixed and Rural environment slopes are represented, with City environment as baseline. Positive values correspond to an advantage compared to growing up in cities. Countries are ranked according to their suburb slope. The slopes of the different non-city environments are highly correlated: Pearson's r(suburb, mixed) = 0.97, $p < 0.001$, r(suburb, rural) = 0.72, $p < 0.001$, r(mixed, rural) = 0.53, $p < 0.001$. The country ranking is very similar to the one with only 2 classes (city / non-city): Spearman's r(non-city, suburb) = 0.85, $p < 0.001$, r(non-city, mixed) = 0.73, $p < 0.001$, r(non-city, rural) = 0.94, $p < 0.001$. P-values are from a t-test testing the hypothesis of no correlation against the alternative hypothesis of a nonzero correlation. **c** - Pairwise differences between random environment slopes shown in panel b, averaged over countries. We show that the average difference in effect size between the city environment and the other 3 environments (city-rural, city-mixed, city-suburb) are around 10 times larger than the difference between the 'non-city' environments (rural-mixed, mixed-surburb, rural-suburb). This supports the approach to cluster together rural, mixed and suburb environments. All error bars correspond to standard errors, n = 397,162 participants.

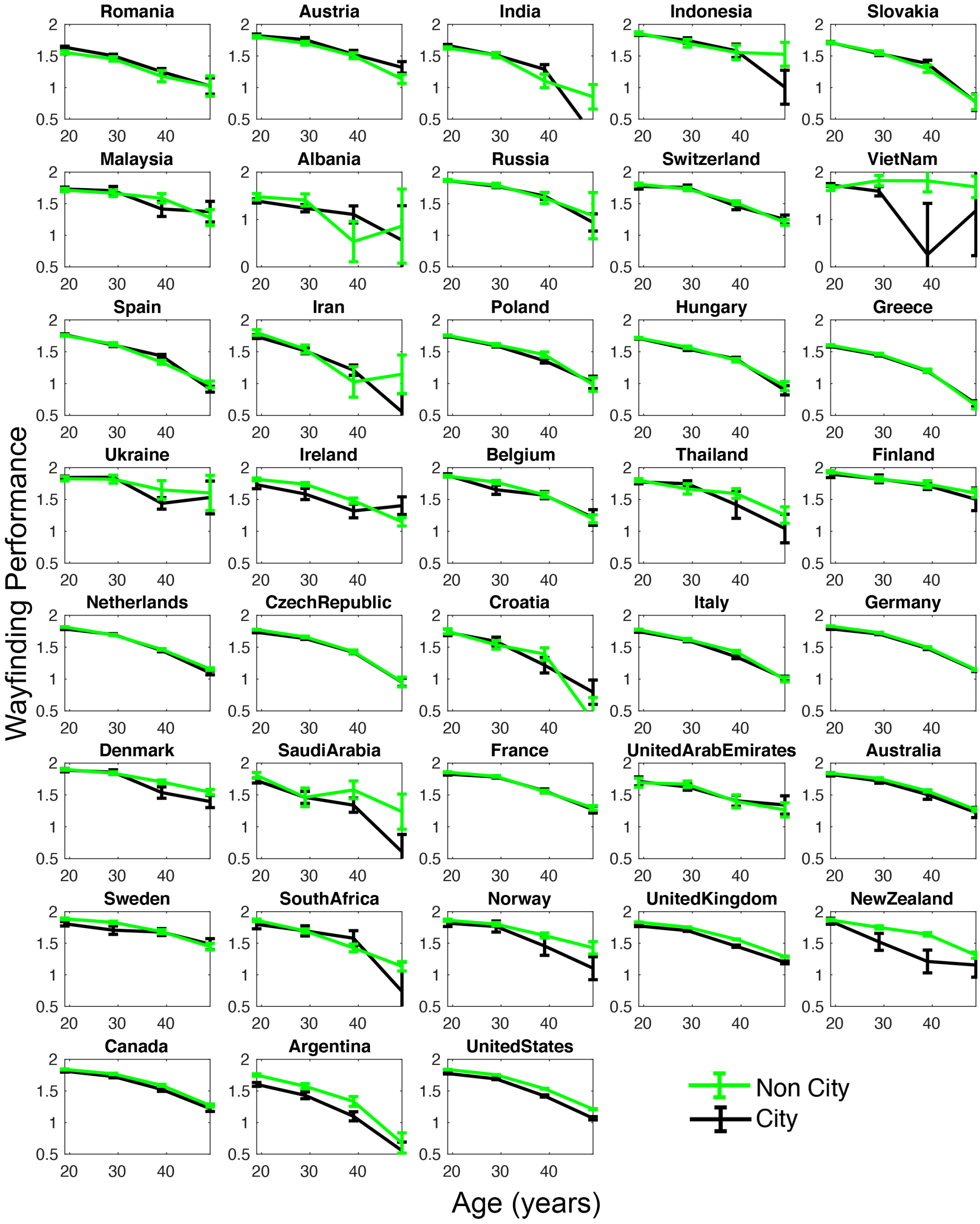


**Extended Data Figure 3. | Wayfinding Performance in city and non-city environments across age, in each country.** Wayfinding Performance is averaged within 10-year windows. Error bars correspond to standard errors and center values correspond to the means. Note that these values correspond to raw Wayfinding Performance, i.e. they have not been corrected for age, gender or education. Note: VietNam and Albania y axis lower bound is 0 to allow display of data points, instead of 0.5 for the rest of the countries. Altogether, we included n = 397,162 participants.

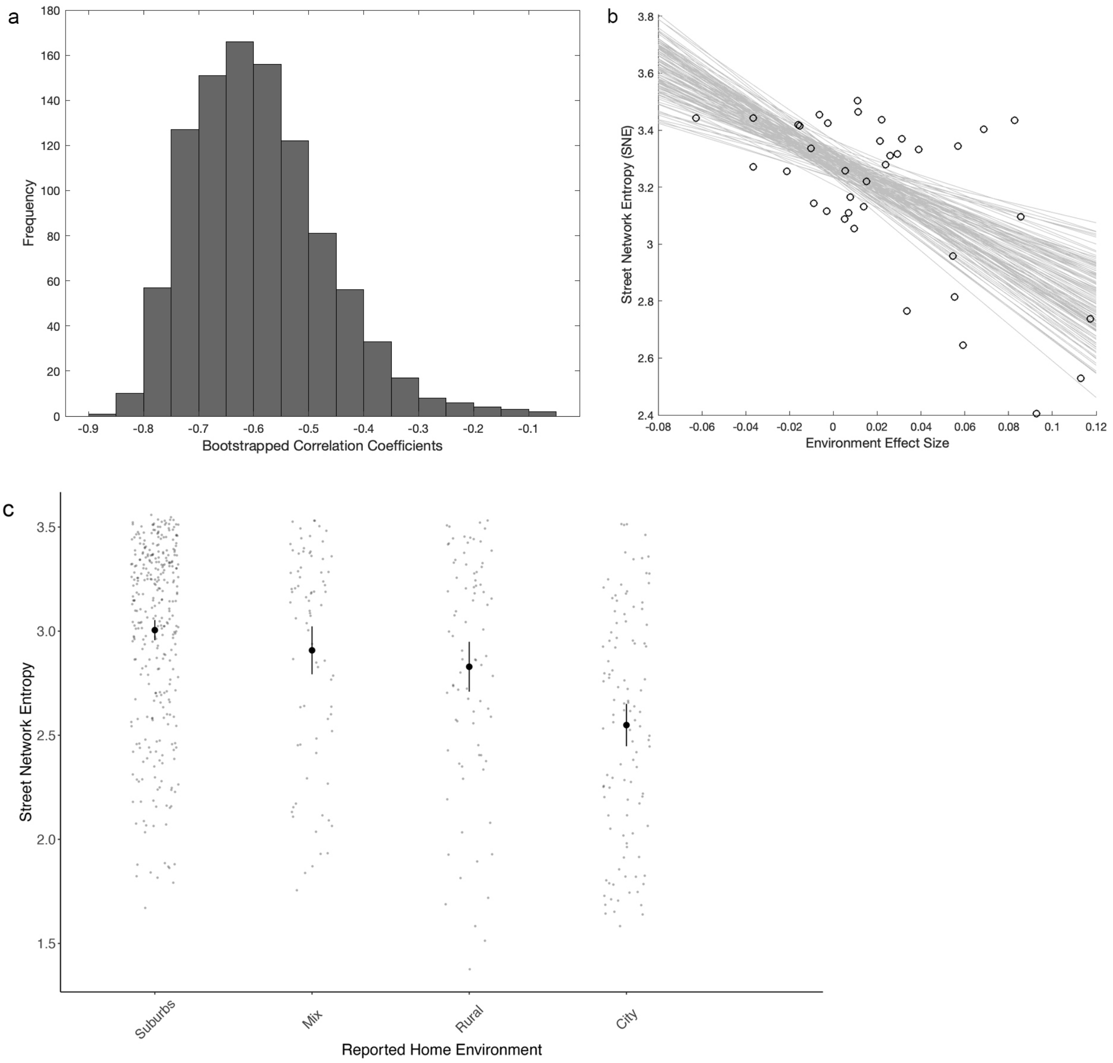


**Extended Data Figure 4. | Estimation of the robustness of the Pearson's correlation between Street Network Entropy (SNE) and environment effect size.** Bootstrapped correlation coefficients computed from 1000 resampling with replacement. **a -** Histogram of the computed correlation coefficients. We obtained r = -0.60, 95% CI = [-0.78 -0.30]. **b -** Regression lines for each sample. **c -** SNE computed at the home addresses of the 599 participants to the follow-up experiment City Hero Quest as a function of the reported type of home environment. The size of the square boxes used to compute the SNE were adjusted for the average street density within each reported environment (see Supplementary Methods). Error bars correspond to standard errors and center values correspond to the means.

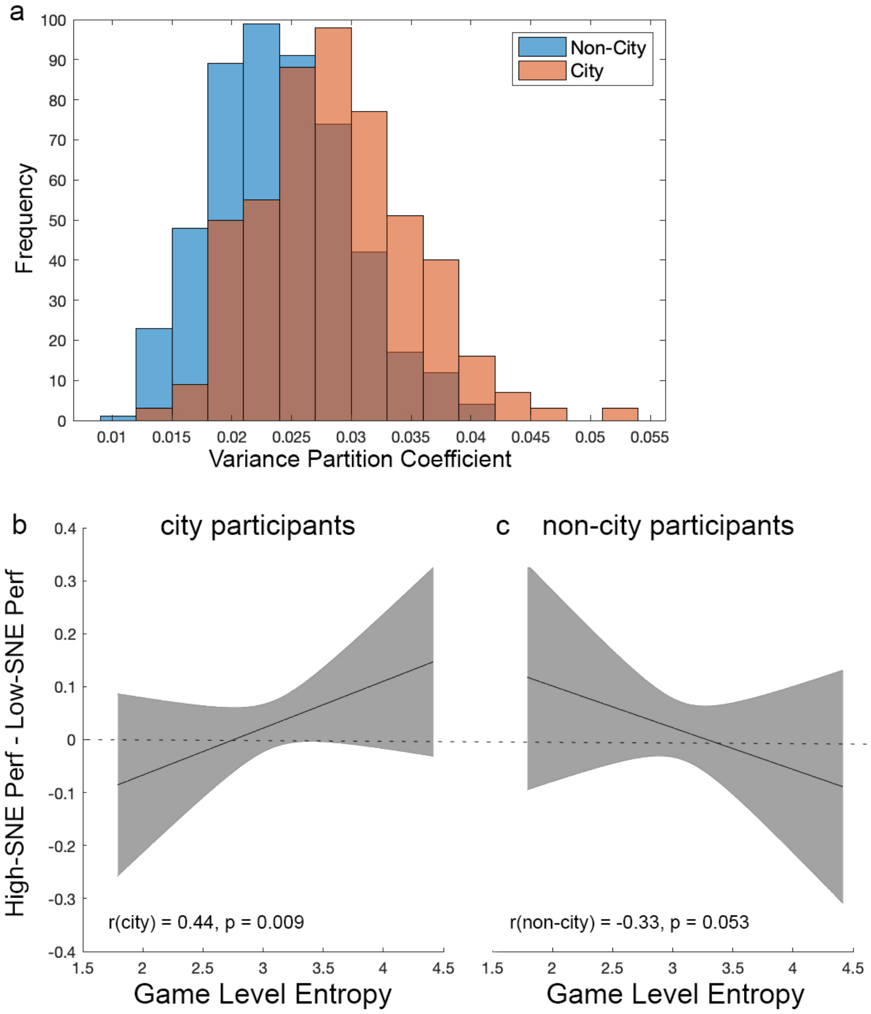


**Extended Data Figure 5.** | **Separation of the effect of country and of game level in city and non-city environments. a -** Bootstrapped distribution of the Variance Partition Coefficient (500 iterations) of the random effect 'country', for city and non-city participants. **b -** Least square regression line of the difference in Wayfinding Performance between city participants from high-SNE countries and city participants from low-SNE countries, as a function of Level Entropy. The y-axis is $D_{city}$, the difference between the level conditional modes of $LMM_{city\_highSNE}$ and the level conditional modes of $LMM_{city\_lowSNE}$. **c -** Least square regression line of the difference in Wayfinding Performance between non-city participants from high-SNE countries and non-city participants from low-SNE countries, as a function of Level Entropy. The y-axis is $D_{noncity}$, the difference between the level conditional modes of $LMM_{noncity\_highSNE}$ and the level conditional modes of $LMM_{noncity\_lowSNE}$. The difference between b and c correlation slopes was significant (Fisher's z = 3.60, $p < 0.001$, 95% CI for r(city) - r(non-city)= [0.36, 1.10]). See Supplementary Notes for more details. Error bars correspond to 95% confidence intervals.

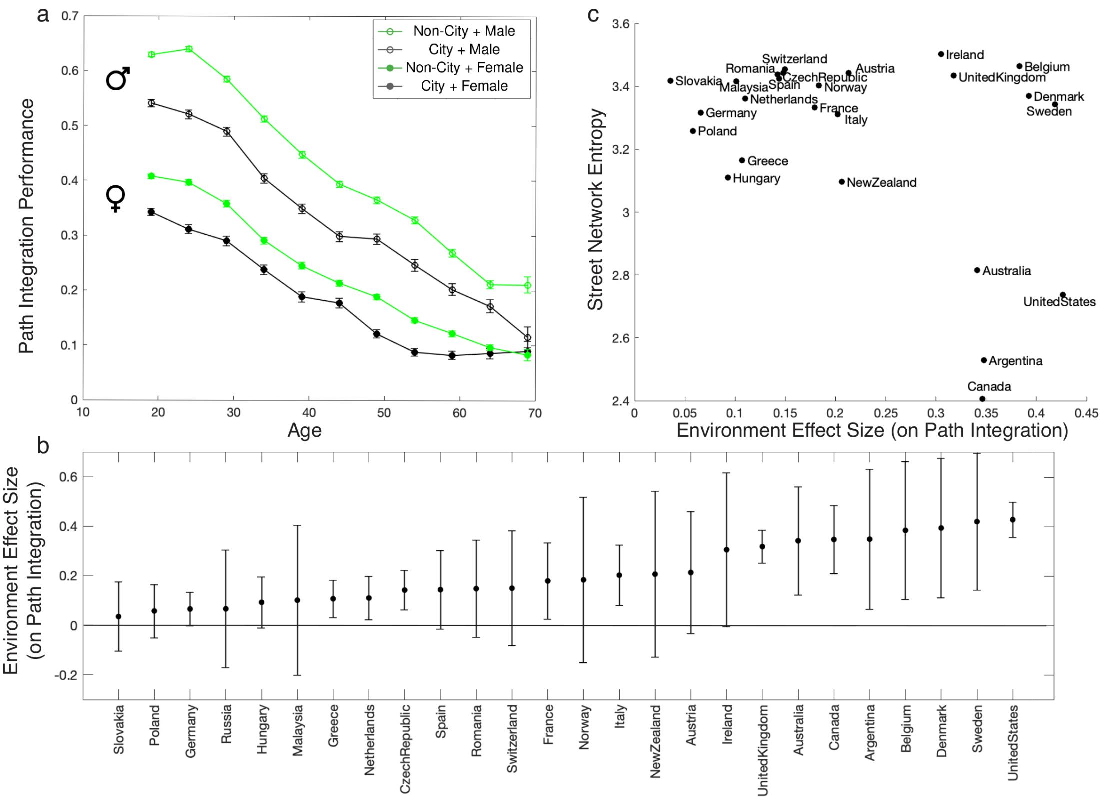


**Extended Data Figure 6. | Association between age, home environment, country, and Path Integration Performance.** a - Path Integration Performance as a function of age for male and female participants who grew up in city and non-city environments. Path Integration Performance is averaged within 5-year windows, center values correspond to the means. b - Difference of the effect of growing up outside cities on Path Integration Performance across countries. We fit a logistic mixed model for Path Integration Performance, with fixed effects for age, gender and education, and random environment slopes clustered by country, see Supplementary Methods. Positive values indicate an advantage for participants raised outside cities. c - Street Network Entropy (SNE) as a function of the environment effect size (random environment slope) in each country, as in Figure 2b, see Supplementary Methods. All error bars correspond to standard errors, n = 182,122 participants.

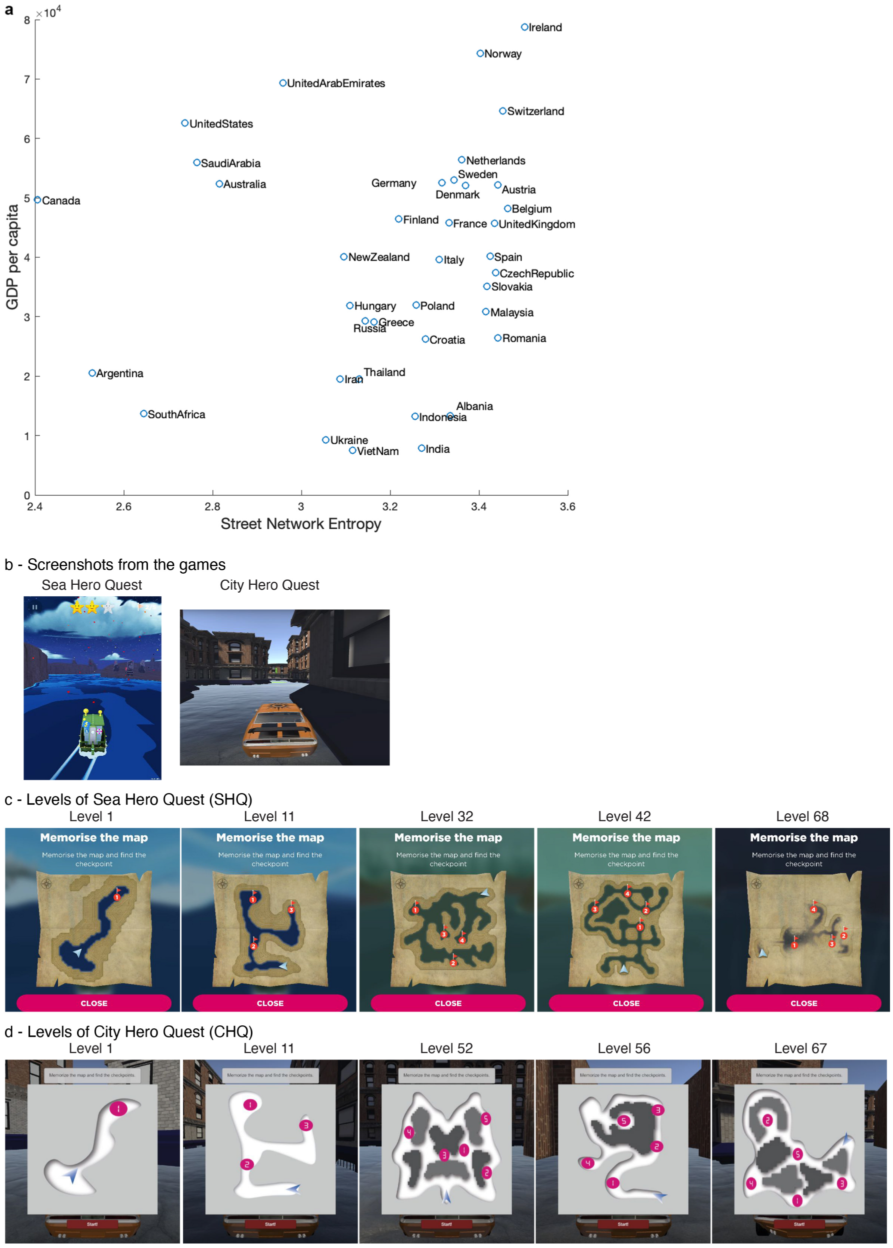


**Extended Data Figure 7.** | **a -** Gross Domestic Product (GDP) per capita as a function of Street Network Entropy. **b -** Screenshot from Sea Hero Quest (SHQ, left) and City Hero Quest (CHQ, right). **c -** Subset of SHQ levels used in the second experiment run on Prolific. **d -** CHQ levels used in the second experiment.

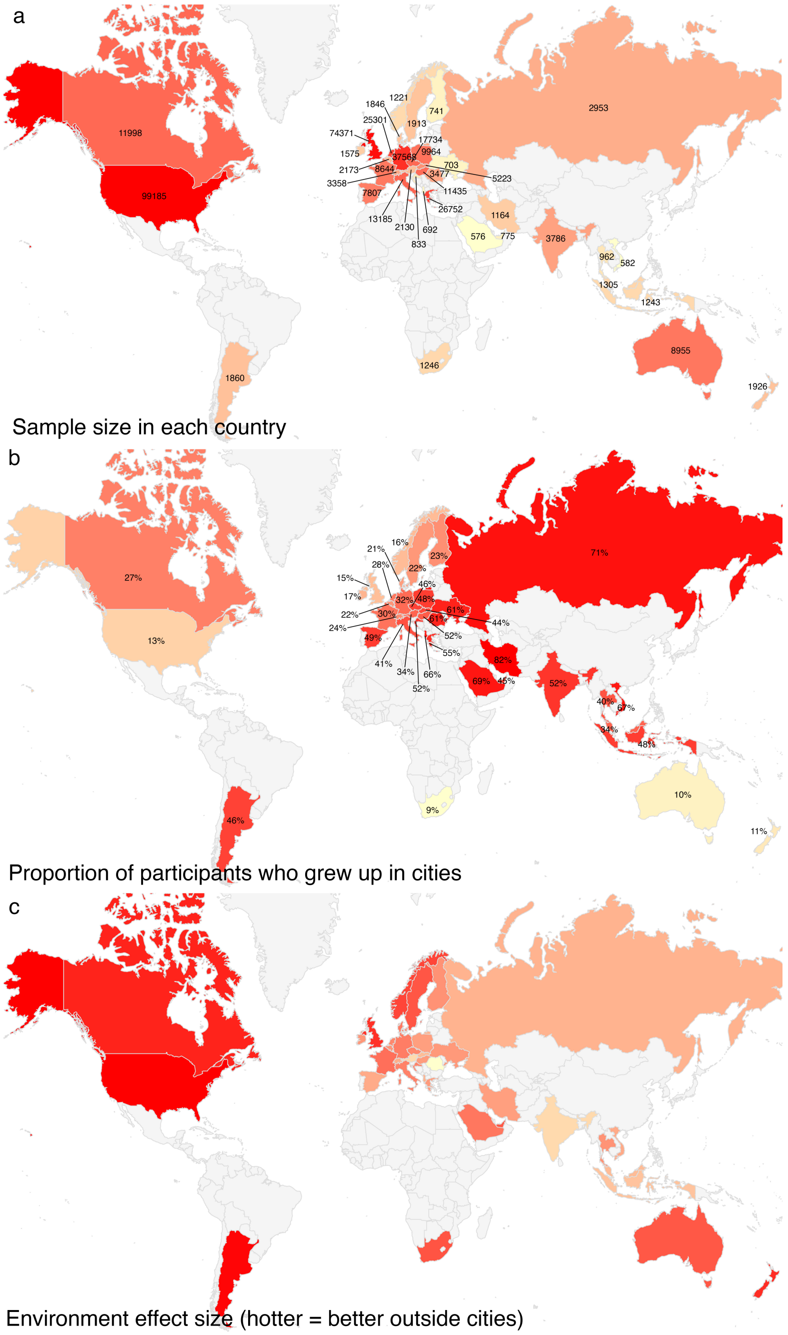


**Extended Data Figure 8.** | **Color-coded world maps.** **a -** Sample size. **b -** Proportion of city participants. **c -** Environment effect size computed from a Linear Mixed Model predicting Wayfinding Performance, with fixed effects for age, gender and education, and random environment slopes clustered by country. The environment effect sizes are the environment slopes clustered by country, identical to the values in Figure 2a.

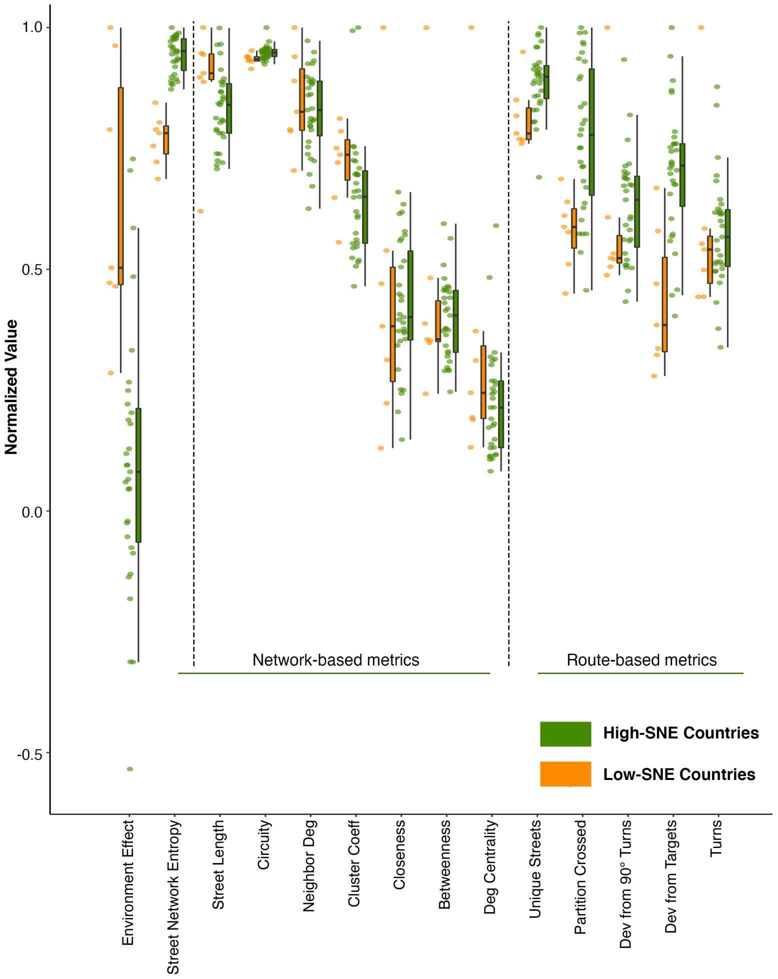


**Extended Data Figure 9. | Environment effect size and city complexity measures in high-SNE and low-SNE countries -** In each of the 380 included cities we computed a range of metrics to quantify different aspects of its complexity. We then took an average of these metrics weighted by the city population to have one value per country. We normalized these values by dividing them by their maximum. **Network-based metrics -** On top of the Street Network Entropy used in this study, we computed other graph-theoretic measures commonly considered for spatial analysis of cities: average street length, circuity, neighborhood degree, clustering coefficient, closeness centrality, betweenness centrality, and degree centrality. **Route-based metrics -** we simulated 1000 routes in each city, and quantified five key variables derived from each route: number of unique streets, number of transitions in the partitions in street network structure, deviation from regular 90 degree turns at each turn, overall deviation from the target and number of turns above 50 degrees. Individual data points correspond to countries (n=38). In the boxplots, the horizontal bar represents the sample median, the hinges represent the first and third quartiles, and the whiskers extend from the hinges to the largest/lowest value no further than ±1.5 * IQR from the hinge (where IQR is the inter-quartile range).

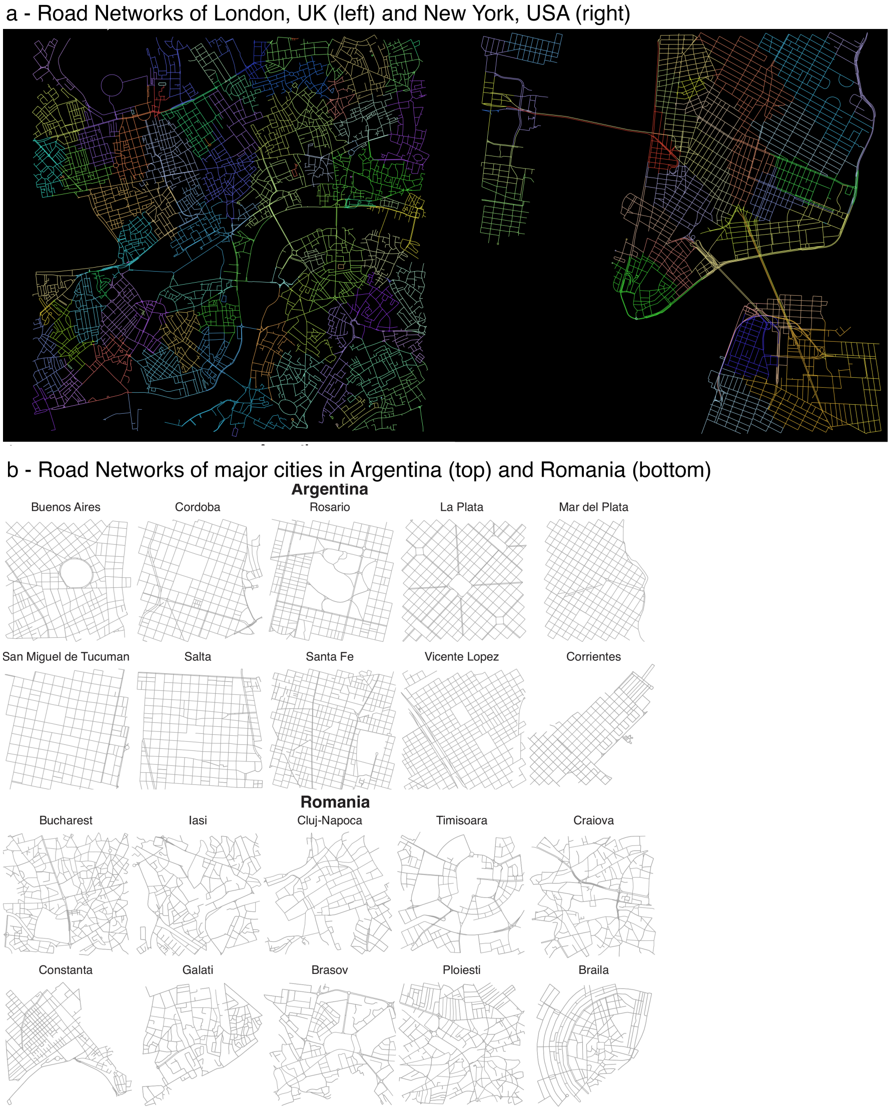


**Extended Data Figure 10.** | **Examples of city street networks - a -** The road networks of New York City (USA, right) and London (UK, left) have been partitioned using the Louvain community detection algorithm on the dual graph, setting edge cost as angular change. The road networks within a 3x3 km$^2$ box around the city centres are represented. **b -** Street network of the 10 biggest cities in terms of population in Argentina and in Romania. We used OSMnx to gather the "drive" Open Street Map network within 1000 $\times$ 1000 m$^2$ boxes around each city centre. The reasons behind these differences are mostly historical. In South America, grid city design is characteristic of Hispanic American colonization [58,59], while disorganized street networks correspond to the typical organic street pattern of old European city cores [60].
